# VPS29 exerts opposing effects on endocytic viral entry

**DOI:** 10.1101/2021.08.06.455441

**Authors:** Daniel Poston, Yiska Weisblum, Alvaro Hobbs, Paul D. Bieniasz

## Abstract

Emerging zoonotic viral pathogens threaten global health and there is an urgent need to discover host and viral determinants influencing infection. We performed a loss-of-function genome-wide CRISPR screen in a human lung cell line using HCoV-OC43, a human betacoronavirus. One candidate gene, VPS29, was required for infection by HCoV-OC43, SARS-CoV-2, other endemic and pandemic threat coronaviruses as well as ebolavirus. However, VPS29 deficiency had no effect on certain other viruses that enter cells via endosomes and had an opposing, enhancing effect on influenza A virus infection. VPS29 deficiency caused changes endosome morphology, and acidity and attenuated the activity of endosomal proteases. These changes in endosome properties caused incoming coronavirus, but not influenza virus particles, to become entrapped therein. Overall, these data show how host regulation of endosome characteristics can influence viral susceptibility and identify a host pathway that could serve as a pharmaceutical target for intervention in zoonotic viral diseases.

## INTRODUCTION

Because viruses rely on host cellular proteins to replicate, an attractive strategy for the next-generation of antiviral therapies is targeted inhibition of human proteins—termed “dependency factors”—that are required for viral replication. Of particular interest are human proteins required by diverse viral lineages, encompassing not only known human pathogens but animal viruses that are of concern for future spillover into human populations. One universal aspect of the viral lifecycle that could be targeted pharmacologically is viral entry. All enveloped viruses require fusion between viral and host cellular membranes for infection (White and Whittaker, 2016). Some enveloped viruses preferentially fuse at the plasma membrane, while others enter cells via endocytosis and fuse in compartments of the endolysosomal system (Grove and Marsh, 2011). Viruses that fuse at the plasma membrane sometimes depend on the expression of cell surface proteases to activate viral fusion proteins, while viruses that enter through endosomes can be highly dependent on endosomal characteristics such as the presence of certain endosomal proteases and/or endosomal pH (Laporte and Naesens, 2017; Marsh and Helenius, 2006).

The specific route of entry can dictate which dependency factors are required for productive infection. For example, the hemagglutinin (HA) of most influenza A virus (IAV) strains must be cleaved by trypsin-like proteases, which primes it for receptor binding and subsequent fusion (Böttcher-Friebertshäuser et al., 2014). Like IAV, the spike protein of coronaviruses must also be processed by proteases in order to enter target cells. However, unlike HA, the spike protein often has two distinct cleavage sites, termed S1/S2 and S2’, that are cleaved during different stages of the virus replication cycle, including biosynthesis (by the Golgi resident furin-like proteases) and during entry (by cell surface TMPRSS2 protease or endosomal cathepsins) (Millet and Whittaker, 2015). Cleavage regulates the liberation of the fusion peptide to enable fusion of the viral envelope with the cellular membranes, allowing infection to proceed. Similarly, the envelope protein of filoviruses, GP, requires two distinct cleavage steps. First, Furin mediated cleavage during exocytosis yields two subunits, GP1 and GP2, which remain linked by disulfide bonds to form the heterodimers that compose the trimeric envelope complex. Following endocytosis, GP1 is further cleaved by endosomal proteases, mainly cathepsins, in a process that removes the cap and the mucin-like domain to enable binding of GP1 to its endosomal receptor, Niemann-Pick C1 (NPC1)(Volchkov and Klenk, 2018).

In this century alone, four emerging zoonotic respiratory pathogens—SARS-Coronavirus (CoV), MERS-CoV, H1N1 influenza A virus (IAV), and SARS-CoV-2—have caused significant morbidity and mortality. Of these, SARS-CoV, MERS-CoV, and SARS-CoV-2 are all enveloped, positive-stranded RNA viruses in the genus betacoronavirus (Coronaviridae Study Group of the International Committee on Taxonomy of Viruses, 2020). Four other coronaviruses are known to infect humans; Human CoV (HCoV)-OC43 and HCoV-HKU1 are members of the betacoronavirus genus (Killerby et al., 2018), while HCoV-229E and HCoV-NL63 are members of the alphacoronavirus genus. Each generally causes only mild illness. To identify coronavirus dependency factors, we performed a genome-wide loss-of-function CRISPR screen using HCoV-OC43 in a human lung cell line, and focused on candidate hits that are required by diverse Coronaviridae. We identified one such factor, VPS29, that is broadly required by both human and animal CoVs. VPS29 is a component of both retromer (VPS26/VPS29/VPS35) and retriever (DSCR3/VPS29/C16orf62), two distinct but related complexes that, together with the CCDC22/CCDC93/COMMD (CCC) complex, mediate endosome-to-plasma-membrane and endosome-to-TGN recycling of transmembrane cargo (Baños-Mateos et al., 2019; McNally et al., 2017; Phillips-Krawczak et al., 2015; Singla et al., 2019). We show that loss of VPS29 impairs CoV infection, and also causes failure of ebolavirus infection. In stark contrast, we show that VPS29 deficiency facilitates IAV infection. We further show that VPS29 deficiency causes profound changes in endosomal properties, including alteration of morphology, acidity and proteolytic activity that differentially impact the egress of viruses from endosomes.

## RESULTS

### A genome wide screen reveals HCoV-OC43 dependency factors

To identify host proteins required for HCoV-OC43 infection, we performed a genome-wide CRISPR screen in the A549 lung adenocarcinoma cell line. Briefly, A549 cells were transduced with the Brunello sgRNA library (Doench et al., 2016; Sanson et al., 2018) at a low MOI (0.3) and high coverage (500X) to generate a population of cells each harboring a single sgRNA. After selection to remove untransduced cells, A549-Brunello cells were infected with HCoV-OC43 at an MOI of 0.1 and incubated for 1 week to allow viral-induced cell death to occur (Figure 1A). Enrichment of sgRNA sequences in the surviving cells—i.e. those putatively lacking a dependency factor—was assessed using MAGeCK (Li et al., 2014).

**Figure 1:**
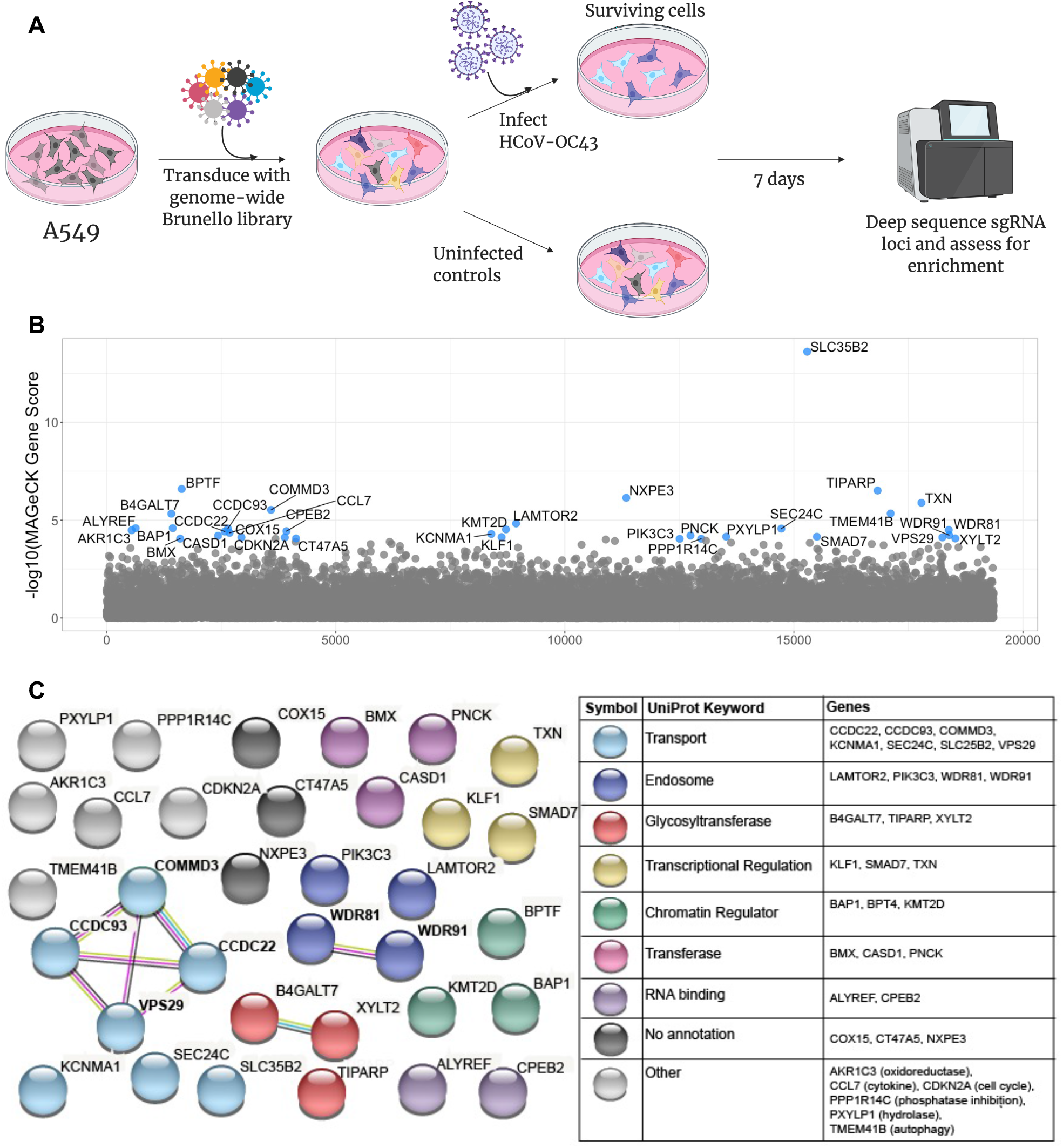
A CRISPR screen reveals genes influencing HCoV-OC43 susceptibility. (A) Schematic of screening setup (B) Screen results, where the x-axis corresponds to each unique gene in the library (labeled randomly from 1 to 19,114) and the y-axis denotes the -log_10_ MAGeCK gene score. All genes scoring higher than the best-scoring non-targeting control pseudogene are labeled in blue. The screen was performed in three independent replicates (C) string-db analysis and UniProt annotation of gene hits. Sphere colors correspond to UniProt keywords and connecting lines indicate strength of evidence underlying gene-gene interactions (pink: experimentally-determined interaction; blue: annotated interaction in curated databases; gray: evidence of co-expression; yellow: text-mining).

We identified 34 candidate dependency factors, defined as genes scoring higher than the highest scored non-targeting control (Figure 1B). As a positive control, we identified CASD1, the enzyme responsible for the generation 9-O-acetylated sialic acids, which serve as the receptor for HCoV-OC43 (Schwegmann-Wessels and Herrler, 2006). Consistent with several other genome wide screens for viral dependency factors, we identified multiple genes (SLC35B2, XYLT2, and B4GALT7) involved in heparan sulfate biosynthesis, implying that heparan sulfate is an attachment factor for HCoV-OC43 (Gao et al., 2019; Luteijn et al., 2019; Milewska et al., 2014; Park et al., 2017; Schneider et al., 2020).

To further classify gene hits (Figure 1C), we performed a functional enrichment analysis using string-db followed by annotation with UniProt keywords (Szklarczyk et al., 2019; UniProt Consortium, 2019). Many of the hits were associated with intracellular transport or endosome activity including VPS29, the **C**CDC22/**C**CDC93/**C**OMMD3 (CCC) complex, and the WDR81/91 complex, suggesting a requirement for these functions in HCoV-OC43 infection. Additionally, we identified PIK3C3, which generates phosphatidylinositol 3-phosphate (PI(3)P), a phospholipid required for the recruitment of retromer to endosomes (Burda et al., 2002). Some of the genes identified by our screen were also recently reported in CRISPR screens utilizing SARS-CoV-2, implying that they are broadly required for coronavirus infection (Daniloski et al., 2020; Zhu et al., 2021).

### Requirement for candidate host factors is both cell type and virus dependent

We next investigated whether the VPS29/CCC complex and the WDR81/91 were required for infection by a diverse panel of respiratory viruses, including coronaviruses. In addition to HCoV-OC43, we tested additional seasonal HCoVs (HCoV-NL63 and HCoV-229E), rVSV/SARS-CoV-2, a chimeric vesicular stomatitis virus encoding the SARS-CoV-2 Spike protein, as well as other pathogenic respiratory viruses: IAV, adenovirus, and respiratory syncytial virus (RSV). We used CRISPR/Cas9 to generate individual cell lines lacking each gene of interest and confirmed knock-out (KO), both by sequencing target loci and by western blot analyses (Figure S1A). Importantly, KO of these genes did not affect cellular viability or proliferation. Because viral dependency factors identified via CRISPR screening might be required in a cell-type specific manner, we evaluated the requirement of these genes for infection in multiple cell lines expressing ACE2 (the receptor for both SARS-CoV-2 and HCoV-NL63): A549-ACE2, HT1080-ACE2, and 293T-ACE2.

Given their function in endosomal trafficking, we hypothesized that these hits would most likely affect viral entry. We therefore performed single-cycle infection assays and quantified infected cells via flow cytometry. There was strong requirement for VPS29/CCC complex as well as WDR81/91 in A549 cells for all CoVs tested (Figure 2A-D). However, the was no requirement these factors in for IAV, adenovirus, or RSV infection of A549 cells (Figure 2E-G). In all other cell lines tested, there was a strong requirement for VPS29 for all coronaviruses but no dependency on VPS29 or the other candidate proteins was found for adenovirus and RSV (Figure 2H-T). Since these viruses all rely on endocytic pathways for viral entry (Krzyzaniak et al., 2013; Lakadamyali et al., 2004; Meier and Greber, 2004), these data indicate that VPS29/CCC and WDR81/91 are specifically required for coronavirus infection, rather than broadly impairing endocytic function. The magnitude of the effect of CCC complex and WDR81/91 knockout on CoV infection was different in different cell lines. For example, KO of the CCC complex or WDR81/91 had a blunted effect on CoV infection in HT1080-ACE2 cells (Figure 2H-K). Moreover, in 293T-ACE2 cells, KO of the CCC complex inhibited HCoV-OC43 but not HCoV-NL63 or rVSV/SARS-CoV-2 infection, while WDR81/91 knockout impaired infection for all three viruses (Figure 2O-Q).

**Figure 2:**
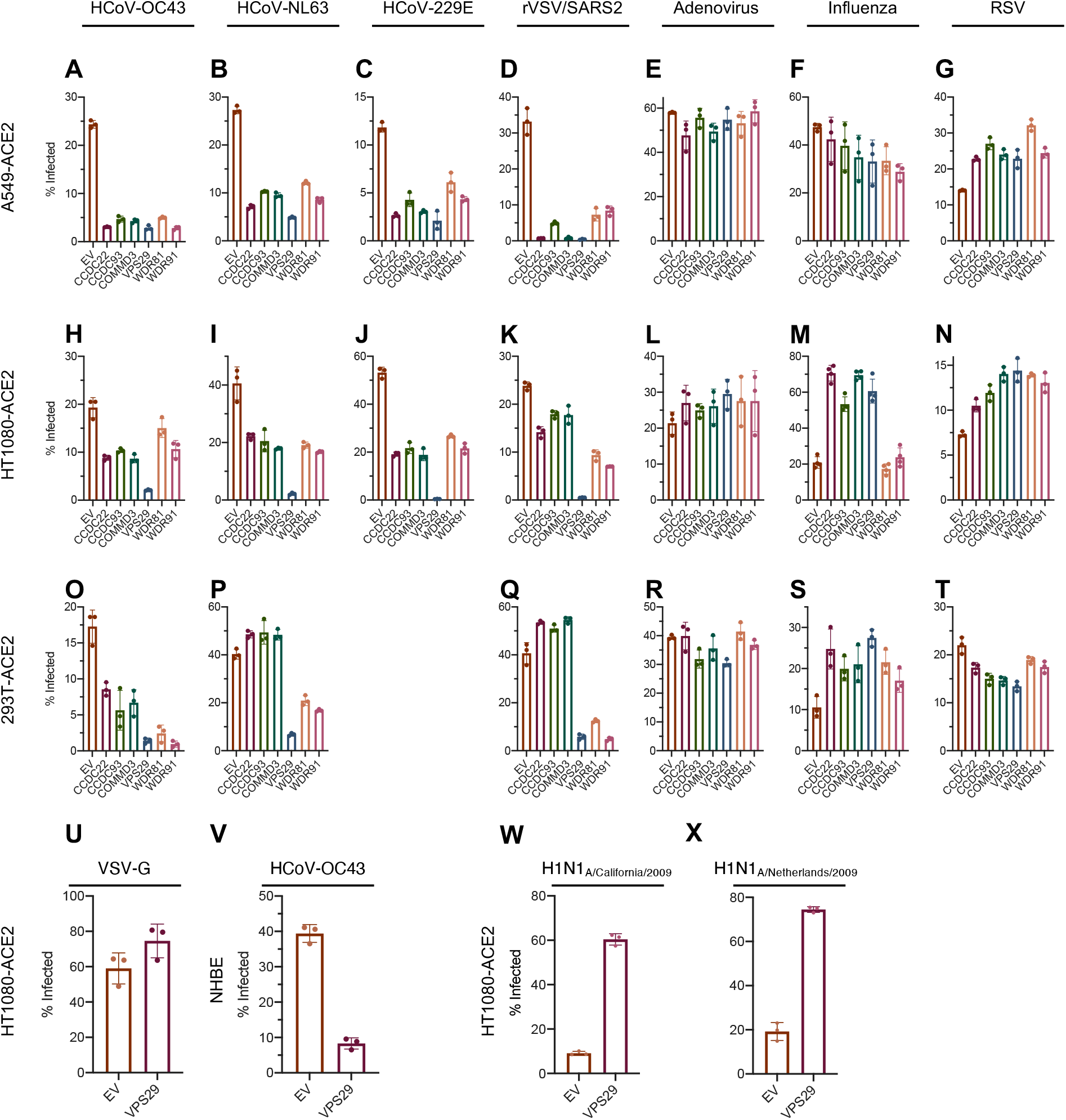
Requirement for identified host proteins is cell type and virus dependent. (A-X) Cells were infected with the indicated viruses at an MOI of 0.3. At 24 hours post infection, cells were stained, and the percent infected cells was determined by flow cytometry. (A-G): A549-ACE2, (H-N, U, W-X): HT1080-ACE2, (O-T): 293T-ACE2, (V): NHBE. X-axis indicates gene knockout, EV: empty vector. Mean (bar graph) of three replicates (dots). Error bars indicate SD. Data shown is a representative of at least two independent experiments.

We found that VSV infection was unaffected by VPS29 KO (Figure 2U). Because the sole difference between rVSV/SARS-CoV-2 and VSV itself is that rVSV/SARS-CoV-2 enters cells using the SARS-CoV-2 spike protein in lieu of VSV-G, these data suggest that it is the entry pathway that imposes the requirement for VPS29. Given the strong requirement for VPS29 by all tested HCoVs, in all cell lines tested, we sought to further confirm the relevance of VPS29 to HCoV infection. To do so, we used CRISPR/Cas9 to KO VPS29 in normal human bronchial epithelial (NHBE) primary lung cells. Loss of VPS29 strongly inhibited HCoV-OC43 infection in NHBE cells (Figure 2V), suggesting that VPS29 is important for HCoV infection of physiologically relevant cells.

In contrast to effects on coronavirus infection, we observed precisely the opposite effect of VPS29 or CCC complex deficiency on IAV infection in HT1080-ACE2 and 293T-ACE2 cells. That is, KO of VPS29 or CCC complex components enhanced IAV infection (Figure 2M,S) while WDR81/91 KO had no effect. To confirm the phenotype observed using the IAV strain A/WSN/33, we analyzed two separate strains of 2009 pandemic H1N1 IAV; A/Netherlands/602/2009 (H1N1)pdm09 (H1N1_2009 Netherlands_) and A/California/04/2009 (H1N1)pdm09 (H1N1_2009 California_). We found that the ability of VPS29 KO to enhance IAV entry was conserved in the pandemic IAV strains (Figure 2W,X). That the same set of endocytic factors could promote infection of coronaviruses while antagonizing IAV infection indicates endosome-based viral entry pathways are influenced by specific sets of host proteins that can facilitate or restrict viral entry.

### VPS29-associated proteins facilitate CoV infection and hinder IAV infection

Because of the opposing effects of VPS29 on HCoV and IAV infection, we elected to examine this protein in more detail—specifically in HT1080 cells, where VPS29 KO strongly suppressed CoV infection and facilitated IAV infection. VPS29 can participate in multiple different protein complexes with distinct roles in normal cell biology (Baños-Mateos et al., 2019). Thus, in order to clarify which VPS29 interacting proteins, if any, are important for facilitating CoV infection and inhibiting IAV infection, we performed a focused siRNA screen targeting VPS29 interacting proteins and assessed impact of knockdown (KD) on HCoV and IAV infection.

Knockdown of VPS26A, VPS29, VPS35, or RAB7A each impaired HCoV-OC43, HCoV-NL63, HCoV-229E, and rVSV/SARS-CoV-2 infection (Figure 3A-D). These data strongly suggest that the participation of VPS29 in the Retromer complex (VPS26A/VPS29/VPS35), which is recruited to endosomes via Rab7A, is the means by which it facilitates CoV infection (Rojas et al., 2008). Interestingly, KD of DSCR3 and C16orf62, which play analogous roles to VPS26 and VPS35 and form the Retriever complex (McNally et al., 2017), inhibited HCoV-OC43 infection but not HCoV-NL63, HCoV-229E, or rVSV/SARS-CoV-2 infection.

**Figure 3:**
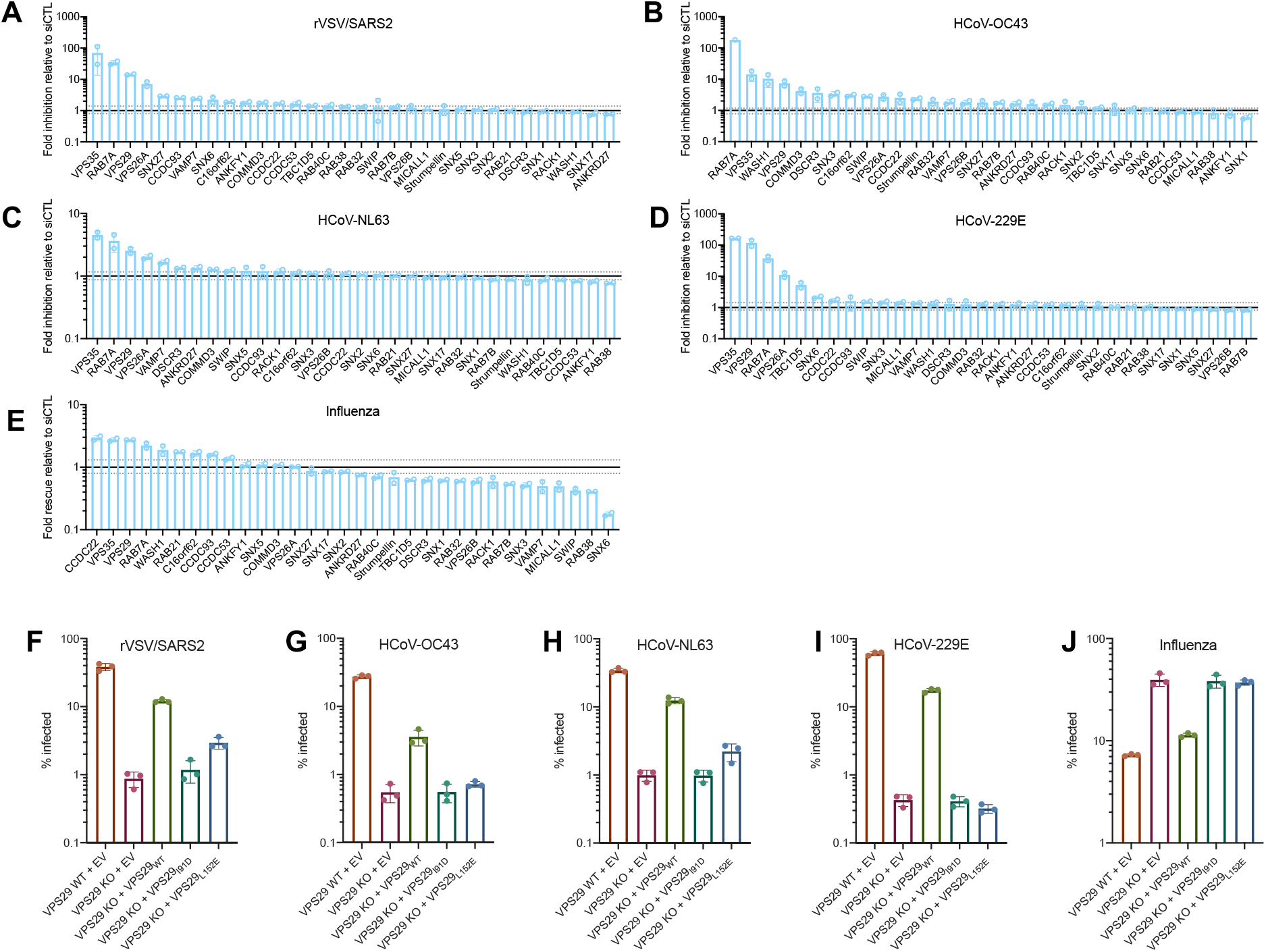
Effect of VPS29 KO on HCoV and IAV infection is primarily driven by loss of Retromer/WASH complex function. (A-E) HT1080 cells were transfected with a focused siRNA library targeting VPS29-interacting proteins. Two days after transfection, cells were infected with (A) rVSV/SARS-CoV-2, (B) HCoV-OC43, (C) HCoV-NL63, (D) HCoV-229E and (E) IAV at an MOI of 0.3. At 24 hours post infection, cells were stained, and the percent infected cells was determined by flow cytometry. Plotted are levels of inhibition (for HCoVs and rVSV/SARS-CoV-2) or increase (for IAV) in siRNA KD cells, relative to siRNA non-targeting control. Fold change values were calculated by comparing levels of infection in KD cells to the average of 4 separate pools of non-targeting siRNA controls. Solid black line marks fold change of 1. The dashed lines mark the highest and lowest fold changes of non-targeting siRNA controls from the average. (F-J) WT and VPS29 mutants were reconstituted in VPS29 KO cells. Cells were infected with (F) rVSV/SARS-CoV-2, (G) HCoV-OC43, (H) HCoV-NL63, (I) HCoV-229E, or (J) IAV. At 24 hours post infection, cells were stained, and the percent infected cells was determined by flow cytometry. Mean (bar graph) of three replicates (dots). Error bars indicate SD. Data shown is a representative of two independent experiments.

IAV infection was enhanced by KD of an overlapping set of VPS29 associated proteins, specifically CCDC22, VPS35, VPS29, RAB7A, WASH1, and RAB21 (Figure 3E). WASH1 is a member of the WASH complex, which facilitates formation of actin patches on endosomes, interacts with and is critical for some protein-sorting functions of retromer (Seaman et al., 2013). RAB21 is a known effector of the WASH complex (Del Olmo et al., 2019). These data thus suggest that the enhancement of IAV infection in VPS29 KO cells is due to the absence of an intact Retromer/WASH complex. While KO or KD of VPS29 facilitates IAV infection, KD of some VPS29-interacting proteins impaired IAV infection. For example, KD of SNX6 impaired IAV infection > 5-fold. However, KD of SNX6 did not affect HCoV infection, indicating that the inhibition of IAV infection is not simply due to global impairment of endosomal function due to SNX6 KD.

### The ability of VPS29 to facilitate CoV infection and inhibit IAV infection depends on interaction with retromer components and regulators

In an orthogonal approach to investigate the role of the Retromer complex in facilitating CoV infection and hindering IAV infection, we generated HT1080 VPS29 KO single cell clones (SCCs) and reconstituted with wildtype (WT) and mutant forms of VPS29. One VPS29 mutant (I91D) does not interact with the Retromer component VPS35, while the other (L152E) does not interact with TBC1D5, a RAB7A GTPase-activating protein that is critical for endosomal recycling of known retromer cargoes (Collins et al., 2005; Harbour et al., 2010; Jia et al., 2016). In agreement with our previous data, CoV infection was inhibited and IAV infection was enhanced in the VPS29 KO SCC (Figure 3F-J). Normal HT1080 cell susceptibility was substantially restored upon reconstitution with a construct expressing a WT, sgRNA-resistant VPS29 (Figure S1 B,C). However, reconstitution with a construct expressing VPS29_I91D_ or VPS29_L152E_ did not reverse the effects of VPS29 KO on HCoV or IAV infection (Figure 3F-J). Overall, these data confirm that loss of the Retromer complex function is the major means by which VPS29 KO affects CoV and IAV infection.

### VPS29 deficiency results in enlarged, deacidified endosomes

To elucidate the impact of VPS29 on viral infection we next investigated the impact of VPS29 KO on normal endosomal function. We labeled endosomes in living cells using a construct containing two FYVE domains fused to mScarlet (2XFYVE-mSCAR), which binds to PI(3)P that is enriched on endosome membranes (Gillooly et al., 2000). Thereafter we treated cells with Dextran labeled with pH-sensitive (pHrodo Green or pHrodo Red) or pH-insensitive (Alexa Fluor (AF)-488) fluorophores to visualize endocytic cargo uptake, as well as the pH status of these endosomes.

Unlike parental HT1080 cells, VPS29 KO cells displayed a prominent subset of enlarged PI(3)P-positive endosomes. These enlarged endosomes were deacidified, as evident from decreased pHrodo Green Dextran signal compared to endosomes in unmanipulated cells, or other smaller endosomes in VPS29 KO cells (Figure 4A,B and S2). Importantly, there was a return to normal endosome phenotype after reconstitution with wildtype VPS29, confirming that this effect is due to VPS29 KO (Figure 4C and S2). The appearance of enlarged, deacidified vesicles was maintained in VPS29 KO cells reconstituted with VPS29_I91D_ or VPS29_L152E_ (Figure 4D,E and S2), suggesting that this phenotype is due to retromer disfunction. Quantification of the pH-sensitive Dextran signal from these images revealed a 3.7-fold decrease in fluorescence intensity in VPS29 KO cells (Figure 4F) that is rescued upon reconstitution with WT VPS29, but not with VPS29_I91D or_ VPS29_L152E._ Importantly, the enlarged endosomes in VPS29 KO cells exhibited equivalent fluorescent intensity to endosomes in normal cells when cells were incubated with pH-insensitive AF-488 Dextran, indicating that while they were deacidified, they were not impaired in cargo loading (Figure 5A, B and S3).

**Figure 4:**
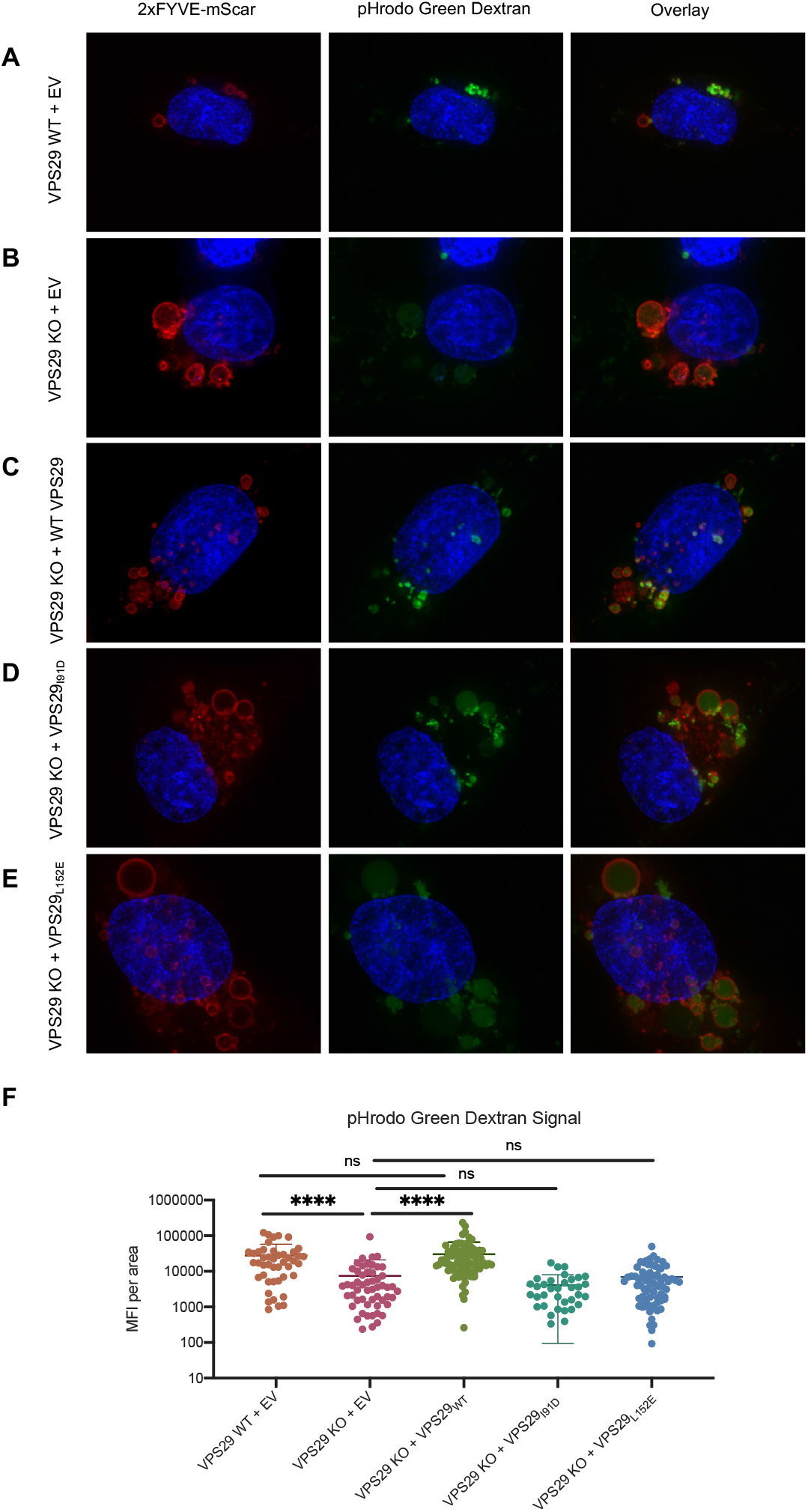
VPS29-KO results in enlarged, deacidified PI(3)P-rich vesicles. Representative images of HT1080 cells transduced with a construct expressing 2xFYVE-mSCAR after incubation with pHrodo Green Dextran for 60 minutes. (A): VPS29 WT + EV expression cassette. (B): VPS29 KO HT1080 + EV expression cassette. (C): VPS29 KO HT1080 reconstituted with WT VPS29. (D): VPS29 KO HT1080 reconstituted with VPS29_I91D_. (E): VPS29 KO HT1080 reconstituted with VPS29_L152E_. EV: empty vector. (F): Quantification of Mean Fluorescence Intensity (MFI) of pHrodo Green Dextran signal inside of 2x-FYVE labeled endosomes from n=4 independent images (images in A-E, as well as the additional representative images depicted in Supplemental Figure S2). Error bars indicate SD. Statistical test: Student’s T test.

**Figure 5:**
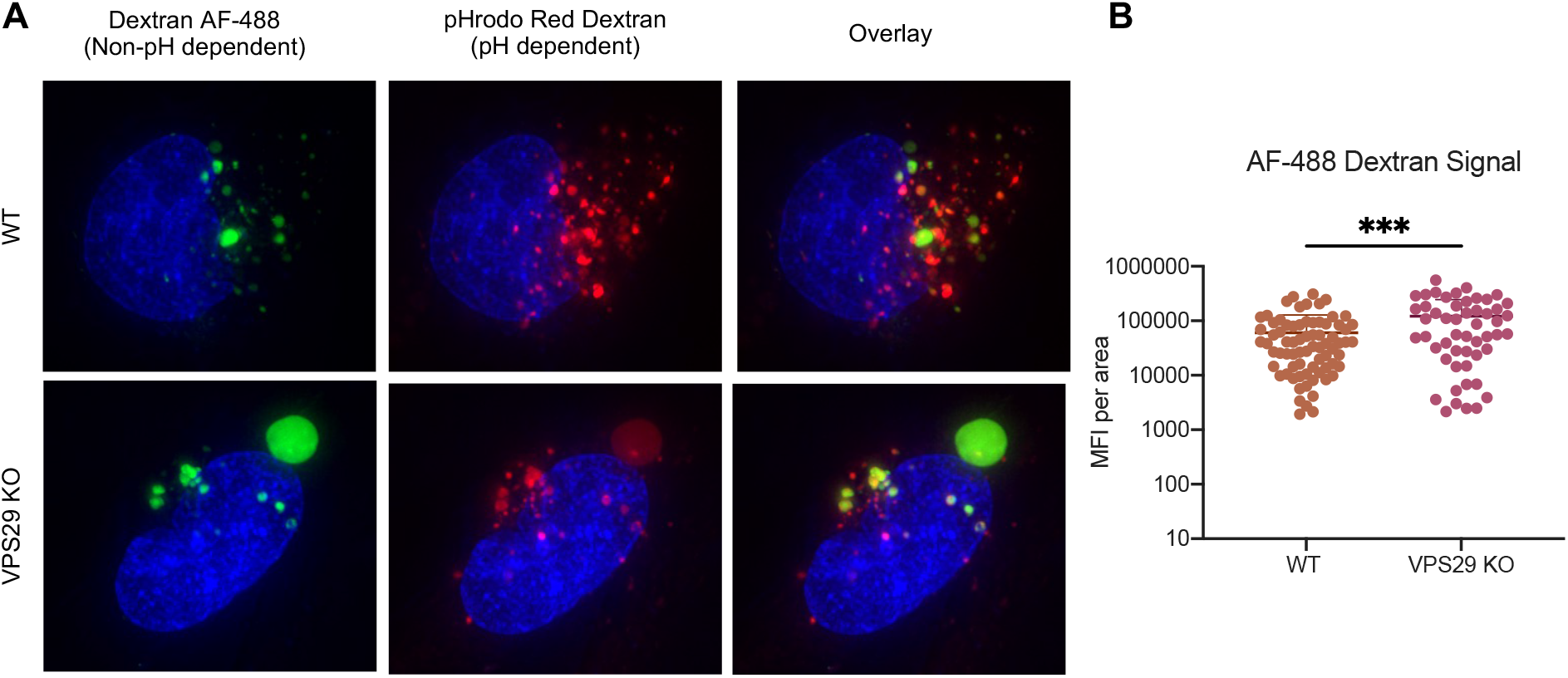
Enlarged, deacidified vesicles in VPS29-KO cells are not impaired for cargo loading. (A): Representative images of HT1080 cells incubated with Dextran AF-488 (non-pH dependent) and pHrodo Red Dextran (pH dependent) for 60 minutes. (B): Quantification of Mean Fluorescence Intensity (MFI) of AF-488 Dextran Signal inside vesicles in WT and VPS29 KO cells from n=3 independent images (images in A, as well as the additional representative images in Supplemental Figure S3). Error bars indicate SD. Statistical test: Student’s T test.

### VPS29 KO results in entrapment of rVSV/SARS-CoV-2 in endosomes

Given the above findings, we hypothesized that CoV infection is impaired in VPS29 KO cells due to impediment in spike dependent egress from endosomes. To test this idea, we generated rVSV/SARS-CoV-2_NG-P_, a replication-competent chimeric VSV expressing SARS-CoV-2 Spike protein in lieu of VSV-G, and containing the VSV structural protein P fused to mNeonGreen (NG-P), thus enabling the direct observation of entering viral particles (Schott et al., 2005).

At 60 minutes post infection of parental HT1080 cells few NG-P punctae were evident within 2xFYVE-mSCAR labeled endosomes, suggesting successful egress of most rVSV/SARS-CoV-2_NG-P_ particles (Figure 6A and S4A) and minimal accumulation therein. However, in VPS29 KO cells, enlarged endosomes contained many rVSV/SARS-CoV-2_NG-P_ punctae at 60 min after infection. Likewise, when cells were infected in the presence of labeled Dextran and imaged 60 minutes post infection, we observed a similar phenotype with rVSV/SARS-CoV-2 particles accumulated in enlarged, Dextran-containing vesicles in VPS29 KO cells (Figure 6B and S4B). Overall, these data indicate that the major inhibitory effect of VPS29 KO on CoV infection is the result of failed egress from endosomes.

**Figure 6:**
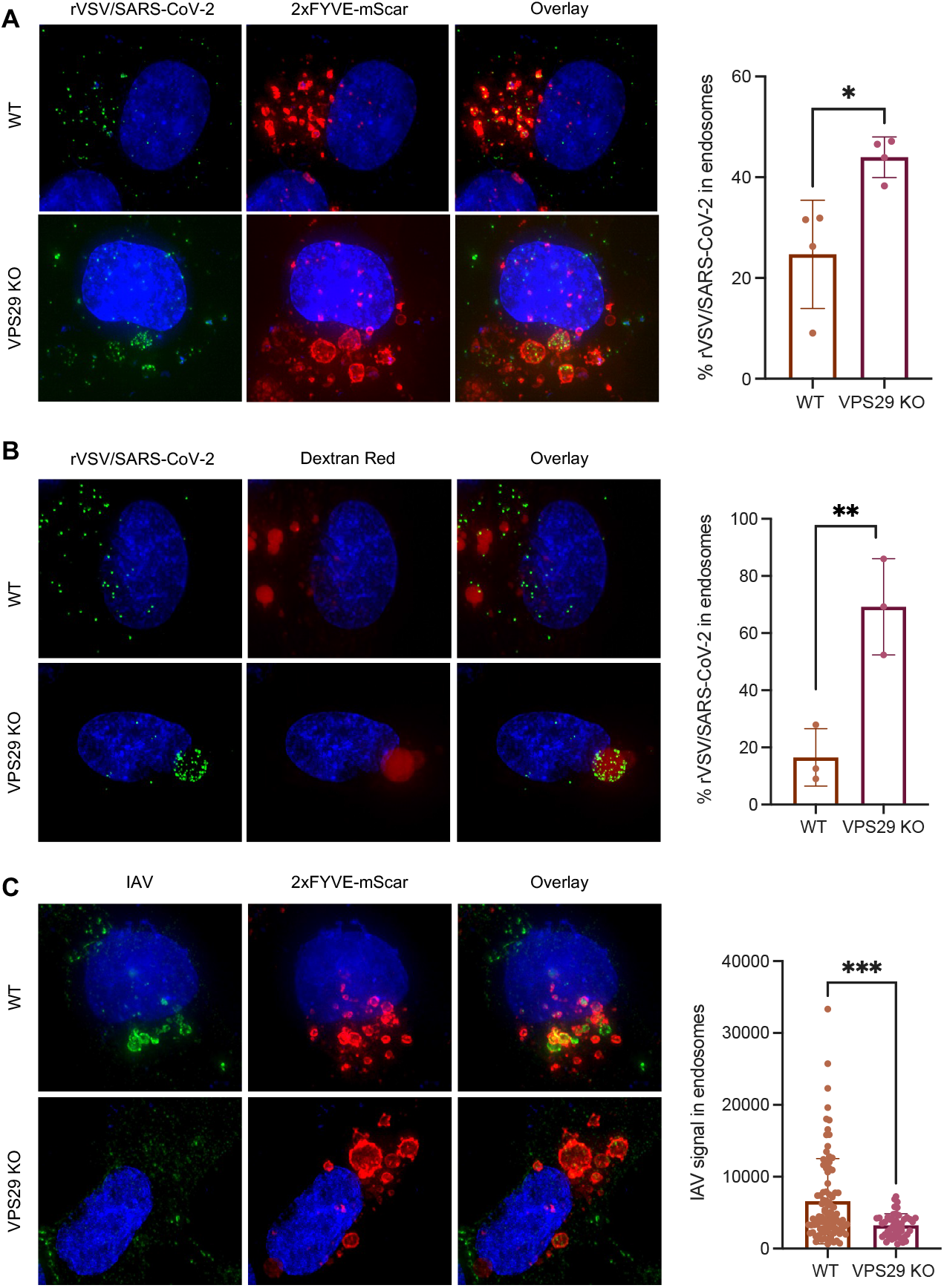
VPS29 KO results in rVSV/SARS-CoV-2 specifically remaining trapped in endosomes. (A) : Representative images of rVSV/SARS-CoV-2_NG-P_ infection in WT and VPS29 KO HT1080 cells. 2xFYVE-mSCAR labeled cells were infected with rVSV/SARS-CoV-2_NG-P_ for 60 minutes. Quantification indicates the percent of rVSV/SARS-CoV-2_NG-P_ punctae inside of 2x-FYVE labeled endosomes from n=4 independent images (images in A, as well as the additional representative images in Supplemental Figure S4A). (B): Representative images of WT and VPS29 KO HT1080 cells incubated for 60 minutes with Dextran Red 10,000 MW and rVSV/SARS-CoV-2_NG-P._ Quantification indicates the percent of rVSV/SARS-CoV-2_NG-P_ punctae inside of Dextran Red labeled endosomes from n=4 independent images (images in B, as well as the additional representative images in Supplemental Figure S4B). (C): Representative images of IAV infection in WT and VPS29 KO HT1080 cells labeled with 2xFYVE-mSCAR. Cells were infected with IAV for 60 minutes then fixed and stained for IAV NP. Quantification indicates the IAV signal (MFI) inside of 2x-FYVE labeled endosomes from n=4 independent images (images in C, as well as the additional representative images in Supplemental Figure S5). Error bars indicate SD. Statistical test: Student’s T test.

Similar experiments in which incoming IAV virions were detected by immunofluorescence 60 min after (Figure 6C and S5) revealed that IAV particles did not accumulate in the enlarged 2xFYVE-mSCAR labeled endosomes in VPS29 KO cells. Thus, the effect of VPS29 KO on rVSV/SARS-CoV-2 was indeed specific. In fact, there was significantly greater association between incoming IAV and 2xFYVE-labeled endosomes in parental HT1080 cells as compared to VPS29 KO cells (Figure 6C and S5), mirroring the opposing effects of VPS KO on HCoV and IAV infection.

We hypothesized that such effect might be due to VPS29-dependent trafficking of antiviral proteins with activity against IAV to endosomes, such as IFITM3. We observed that IFITM3 knockdown enhanced IAV infection of parental HT1080 cells (Figure S6A), in agreement with previous reports (Feeley et al., 2011). However, IFITM3 knockdown augmented IAV infection in VPS29 KO cells (Figure S6A), suggesting that the enhancement of IAV infection in VPS29 KO cells was not the result of loss of IFITM3 activity. Concordantly, IFITM3 localized to 2xFYVE-labeled endosomes in both WT and VPS29 KO cells, and there was no clear difference in localization (Figure S6B). Overall. these finding suggest that enhanced IAV infection in VPS29 KO cells is due to increased egress from endosomes but is not due to altered localization and/or impaired activity of IFITM3.

### Impairment of CoV and ebolavirus infection in VPS29 KO cells due to loss of endosomal cathepsin activity

The aforementioned findings indicate that the reduced susceptibility to HCoV infection in VPS29 KO cells is spike-specific and is the consequence of failed egress from endosomes. We hypothesized that this effect could be due to impaired spike processing by endosomal proteases during entry. We used HIV-1-based pseudotyped viruses to test the susceptibility of various CoV spikes to VPS29 KO and cathepsin inhibition using the drug E64d. As rVSV/SARS-CoV-2 bears a point mutation, R683G, which ablates the polybasic furin cleavage site, we tested pseudotypes bearing WT or R683G mutant spike proteins, as well as spike proteins from SARS-CoV and SARS-like CoV from bats and pangolins, which also do not contain polybasic cleavage sites (Coutard et al., 2020).

Pseudotypes bearing both the WT and R683G mutant SARS-CoV-2 spike proteins were sensitive to VPS29 KO and cathepsin inhibition. However, cathepsin inhibition did not further decrease infection of VPS29 KO cells (Figure 7A). The SARS-CoV-2_R683G_ (Figure7B), SARS-CoV (Figure7C), and the SARS-like bat (Figure 7D) and pangolin viruses (Figure 7E,F) that lack furin cleavage sites were more impacted by VPS29 KO and cathepsin inhibition than WT SARS-CoV-2. Indeed, in several instances, VPS29 KO and/or cathepsin inhibition resulted in undetectable infection by SARS-CoV-2_R683G_, SARS-CoV, and the SARS-like bat/pangolin CoVs. Similarly, infectivity assays utilizing rVSV/SARS-CoV-2 also revealed a dose-dependent inhibition of infectivity upon cathepsin inhibition in parental HT1080, but no impairment of infection upon cathepsin inhibition in VPS29 KO cells (Figure 7G).

**Figure 7:**
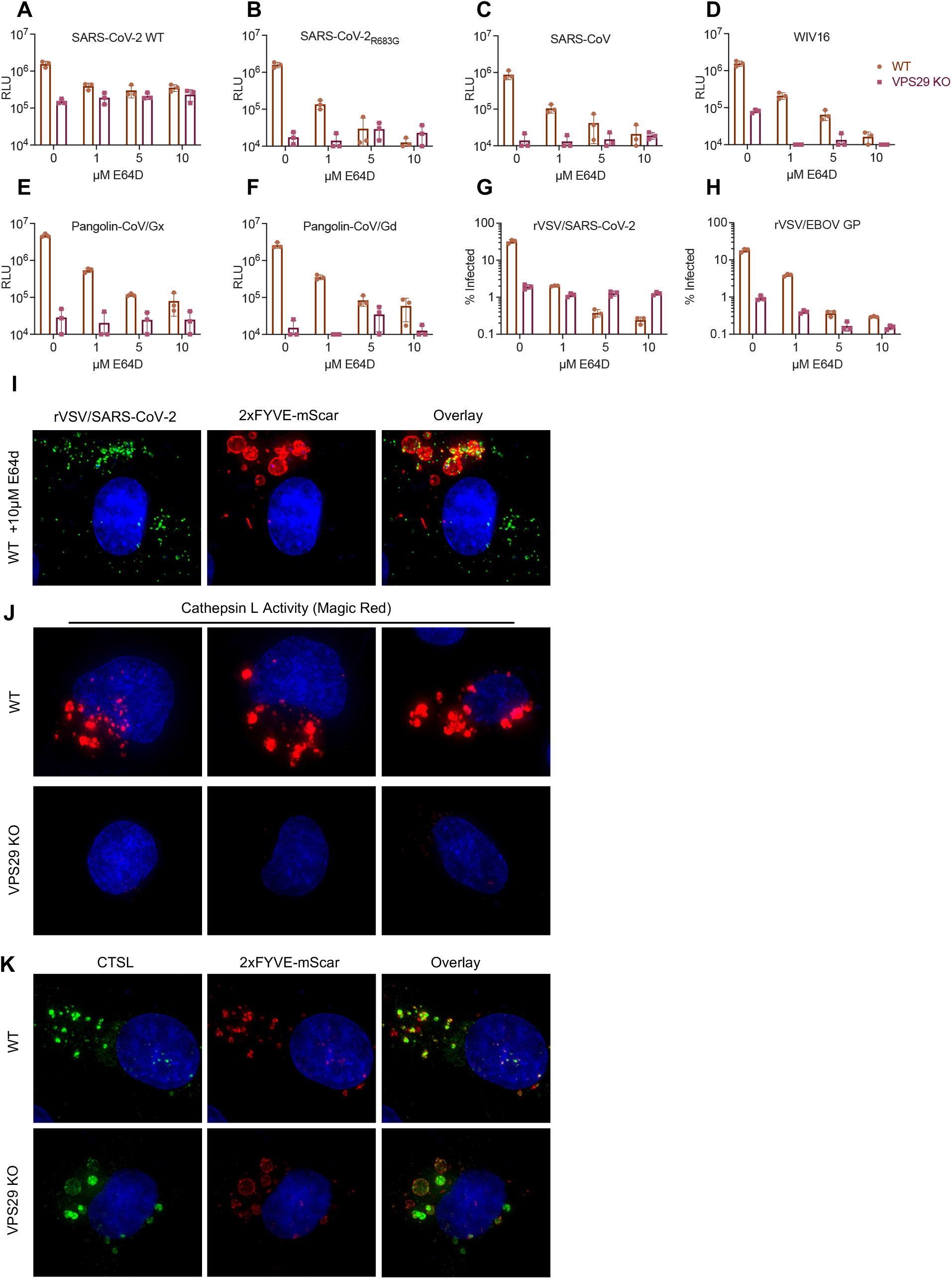
Impairment of CoV infection by VPS29 KO is influenced by presence of polybasic cleavage site and correlates with cathepsin inhibition. (A-F): WT and VPS29 KO HT1080 cells were treated with the indicated concentrations of E64d for 30 minutes before infection with HIV-1 based nano-luciferase reporter viruses pseudotyped with Spike protein of (A): WT SARS-CoV-2, (B): SARS-CoV-2_R683G_, (C): SARS-CoV, (D): WIV16: ,(E): Pangolin-CoV/Gx, (F): Pangolin-CoV/Gd. At 48hpi cells were harvested and nano-luciferase activity was measured. Limit of detection of the HIV-1-based pseudoassay = 10^4^ RLU. (G-H): WT and VPS29 KO HT1080 cells were treated with the indicated concentrations of E64d for 30 minutes before infection with (G): rVSV/SARS-CoV-2 or (H): rVSV/EBOV-GP. At 16 hours post infection, and infected cells enumerated by determined by flow cytometry. Limit of detection of the flow cytometry assay = 0.1 % infection. Mean (bar graph) of three replicates (dots). Error bars indicate SD. Data shown is a representative of at least two independent experiments. (I): Representative images of rVSV/SARS-CoV-2_NG-P_ infection in E64d treated WT HT1080. 2xFYVE-mSCAR labeled cells were treated with E64d for 30 minutes, then infected with rVSV/SARS-CoV-2_NG-P_ for 60 minutes. (J): Representative images of WT and VPS29 KO HT1080 cells following 60-minute incubation with Magic Red Cathepsin L Activity Kit. (K): Representative images of WT and VPS29 KO HT1080 cells stably expressing V5-tagged CTSL and labeled with 2xFYVE-mSCAR.

That there was no further effect of cathepsin inhibition on CoV infection in VPS29 KO cells suggests that the effect of these two manipulations converge on a common pathway in promoting egress from endosomes. We thus hypothesized that VPS29 KO impedes CoV infection by impairing proper processing of spike by cathepsins. If this were indeed the case, then VPS29 KO should impair infection mediated by ebolavirus (EBOV) glycoprotein (GP), which is known to require processing by endosomal cathepsins (Schornberg et al., 2006). To test this, we performed infectivity assays in WT and VPS29 KO cells using a recombinant VSV expressing EBOV GP in lieu of VSV-G (rVSV/EBOV-GP) (Mulherkar et al., 2011). Indeed, we observed a strong inhibition of rVSV/EBOV-GP with both cathepsin inhibition and loss of VPS29 (Figure 7H).

This result suggests that the susceptibility of VPS29 KO is mediated by impaired cathepsin activity.

Consistent with the above conclusion, when parental HT1080 cells were treated with the cathepsin inhibitor E64d, infected with rVSV/SARS-CoV-2 _NG-P_ and examined microscopically, we observed a phenotype similar to that seen in VPS29 KO cells (see Figure 6A). Specifically, substantially more rVSV/SARS-CoV-2_NG-P_ punctae were evident within endosomes, and the endosomes appear enlarged with similar appearance and morphology to those observed in VPS29 KO cells (Figure 7I and S7). To directly test whether VPS29 KO results in impaired endosomal cathepsin activity, we measured endosomal cathepsin activity in WT and VPS29 KO HT1080 cells using a substrate that generates a fluorescent signal upon cleavage by cathepsin L. Indeed, in WT cells, we observed a strong red fluorescence signal in vesicular structures, indicating high levels of cathepsin activity. However, in VPS29 KO cells the red fluorescence signal was nearly absent, indicative of impaired cathepsin activity in VPS29 KO cells (Figure 7J). To determine if the loss of cathepsin L activity was the result of failed trafficking of cathepsins to the endolysosomal system, we performed immunofluorescence studies utilizing tagged cathepsin L in cells with endosomes labeled with 2xFYVE-mSCAR. There was no change in cathepsin L localization to 2xFYVE-mSCAR-positive endosomes in VPS29 KO cells (Figure 7K). These data suggest that the loss of cathepsin activity in VPS29 KO cells is not a result of impaired trafficking of cathepsin itself to endosomes, but rather change endosomal conditions in VPS29 KO cells, such as increased pH, reduces cathepsin activity therein.

## DISCUSSION

While the advent of robust, high-throughput screening modalities has generated a wealth of information regarding host-viral interactions, the underlying mechanism of action for many host proteins implicated by these screens remain incompletely understood. Here, utilizing HCoV-OC43 as a model HCoV, we employed a genome-wide loss-of-function CRISPR screen to identify and characterize factors required for efficient CoV infection. In particular, we show that the retromer subunit protein VPS29 is required for productive infection by diverse CoVs in a variety of cell types. Other genome-wide screens using SARS-CoV-2 have also suggested a role for VPS29 and the CCC as well as RAB7A, which recruits retromer to endosomes (Daniloski et al., 2020; Hoffmann et al., 2021; Wang et al., 2021; Zhu et al., 2021) in HCoV infection

While previous studies have hypothesized that the mechanism whereby VPS29/CCC complex facilitates SARS-CoV-2 infection is by maintaining cell surface expression of viral receptors (Daniloski et al., 2020; Zhu et al., 2021), our findings indicate a different role. Indeed, infection by HCoVs that use three distinct receptors was inhibited by VPS29/CCC KO, while there was no VPS29/CCC requirement for infection by IAV, adenovirus, or RSV—which should otherwise also be dependent on cell-surface expression of their respective receptors. Indeed, HCoV-OC43’s purported receptor (9-O-acetylated sialic acid) is nearly identical to IAV’s receptor (sialic acid). Yet, while HCoV-OC43 infection is highly dependent on VPS29/CCC, IAV infection is either unaffected by these factors (in A549 cells) or hindered by them (in HT1080 and 293T cells).

A possible mechanism that might account for the enhancing effect VPS29 deficiency on IAV infection could be retromer-dependent trafficking of endosomal cargo that antagonizes IAV infection. While this notion is consistent with our finding that showing incoming IAV accumulated in endosomes to a greater extent in normal than in VPS29 KO cells, the proteins that might be responsible for mediating this effect are unknown. Other possibilities, include direct influence of retromer on IAV containing endosomes, In this regard, retromer dependent movement of human papilloma virus (HPV) to the TGN has been reported to involve direct interaction between retromer and the HPV L2 protein (Popa et al., 2015). While loss of retromer/WASH complex facilitated IAV infection in HT1080 cells, loss of other retromer-interacting proteins, such as SNX6 and Rab38, impaired IAV infection. Thus, distinct effector functions of VPS29 may have different infection enhancing and inhibiting properties, with the overall effect depending on viral and/or cell-type specific characteristics. Differences in expression and/or activity of various VPS29 effector proteins may explain why VPS29 KO facilitates IAV infection in HT1080 and 293T cells, but not in A549 cells.

However, VPS29 KO impaired CoV infection in all cells tested, including primary lung cells. Nevertheless, we did observe some differences. Specifically, loss of both retromer and retriever impaired HCoV-OC43 infection, while loss of retriever did not impair HCoV-NL63 or HCoV-229E or rVSV/SARS-CoV-2 infection. These findings suggest that there may be multiple roles for VPS29 in HCoV infection, with some CoVs requiring the effector functions of distinct VPS29-containing complexes. The precise requirement for distinct VPS29 functions could vary with cell type, for example there was a decreased requirement for the CCC complex in HT1080 and 293T cells. Our finding that SARS-CoV-2_R683G_ , SARS-CoV, and bat/pangolin CoVs were all heavily impacted by both VPS29 KO and cathepsin inhibition suggests that these viruses are especially sensitive to endocytic function, in line with recent work demonstrating that mutation of the SARS-CoV-2 polybasic cleavage site drives virions to enter via the endocytic route (Winstone et al., 2021).

Based on our findings, it appears that a key feature of VPS29/retromer KO cells, is elevation of the pH of endolysosomal compartments. This change should impair activation of cathepsins (Jerala et al., 1998), thus impeding endosomal CoV spike as well as EBOV GP processing and egress from endosomes to initiate productive infection, consistent with our observations of incoming virions. In agreement with this model, others have shown VPS35 deficiency results reduced endosomal cathepsin activity (Cui et al., 2019). Here, however, we show that this reduced endosomal activity is likely due to perturbed endolysosomal pH rather than impaired trafficking of cathepsin zymogen in VPS29 KO cells. We also speculate that the perturbations in endosomal pH that accompany VPS29 deficiency underlies increased cellular susceptibility to IAV infection, perhaps by reducing virion exposure to destructive lysosomal proteases, or increasing the duration of exposure to optimal conditions for HA-mediated membrane fusion.

Importantly, our findings suggest that the exploration of cathepsin inhibitors, or other endosomal perturbing agents is a promising target for novel drugs against CoVs, which remain a potentially serious emergent public health threat.

## ACKNOWLEDGMENTS

We thank members of the Bieniasz Laboratory for helpful comments and discussion. This work was supported by grants from National Institute of Allergy and Infectious Diseases, R01AI 50111 and R37AI640003 to (PDB), by a Medical Scientist Training Program grant from the National Institute of General Medical Sciences T32GM007739 to the Weill Cornell/Rockefeller/Sloan Kettering Tri-Institutional MD-PhD Program and by the National Institute of Allergy and Infectious Diseases F30AI157898 (to DP).

## AUTHOR CONTRIBUTIONS

YW, DP, and PDB conceived the study. YW, DP, and AH performed experiments and analyzed the data. DP, YW, and PDB wrote the manuscript.

## RESOURCE AVAILABILITY

### Lead contact

Further information and requests for resources and reagents should be directed to and will be fulfilled by the lead contact, Paul Bieniasz

### Material Availability

Newly generated materials associated with this study are available from the Lead Contact with a completed Materials Transfer Agreement.

### Data and code availability

Sequencing data will be made available upon manuscript acceptance.

## EXPERIMENTAL MODELS AND SUBJECT DETAILS

### Cell culture

HEK-293T (*H. Sapiens;* sex: female), A549 (*H. Sapiens;* sex: male), HT1080 (*H. Sapiens;* sex: male), MDCK (*Canis familiaris*) and Vero cells (*Cercopithecus aethiops*) were obtained from ATCC, and Huh7.5 cells (generously provided by Charles M. Rice) were maintained at 37°C and 5% CO_2_ in Dulbecco’s Modified Eagle Medium (DMEM, Gibco) supplemented with 10% fetal bovine serum. NHBE cells (*H. Sapiens*) were obtained from ATCC (Cat# ATCC PCS-300-010) and maintained at 37°C and 5% CO_2_ in Airway Epithelial Cell Basal Medium (ATCC PCS-300-030) supplemented with Bronchial Epithelial Cell Growth Kit (ATCC PCS-300-040). All cells have been assessed for Mycoplasma contamination.

### Production of viral stocks

HCoV-OC43 (strain: ATCC VR-759) and HCoV-229E (strain: ATCC VR-740) were obtained from Zeptometrix Corporation, and HCoV-NL63 (strain: Amsterdam I) was obtained from the Biodefense and Emerging Infections Research Resources Repository. Viral stocks were generated by propagation on Huh7.5 cells. The IAV strains A/WSN/33 (H1N1), A/Netherlands/602/2009 (H1N1)pdm09 (H1N1_2009 Netherlands_), A/California/04/2009 (H1N1)pdm09 (H1N1_2009 California_) were propagated in MDCK cells. RSV strain A2-line19F expressing the red fluorescent protein monomeric Katushka 2 (mKate2;(Hotard et al., 2012)) was propagated in Vero cells. Adenovirus 5 was purchased from ATCC (VR-1516) and propagated in A549 cells. VSV_IND_(eGFP) was propagated on 293T cells (Whelan et al., 2000). The replication-competent chimeric recombinant vesicular stomatitis virus encoding SARS-CoV-2 S and green fluorescent protein (eGFP), rVSV/SARS-2/GFP_2E1_, has been described previously (Schmidt et al., 2020) and was propagated on 293T-ACE2 cells. rVSV/EBOV-GP was propagated on Vero cells as previously described (Mulherkar et al., 2011).

## METHOD DETAILS

### CRISPR-Cas9 screening

The human genome-wide Brunello Library (Doench et al., 2016) in lentiCRISPRv2 was obtained from Addgene (cat# 73179) and amplified according to depositor’s instructions. Resulting plasmid DNA was validated via NGS sequencing to confirm appropriate coverage and representation (the resulting library contained 0.0% undetected guides and a skew ratio of the top 10% represented guides to the bottom 10% represented guides was 3.94, well below the recommended cutoff of 10 for an “ideal” library (Joung et al., 2017)). To generate lentiviral preparations of the Brunello library, 293T cells (6 × 10^6^ cells per 10 cm dish) were transfected with 6µg lentiCRISPRv2-Brunello, 6µg NL-gagpol, and 1.2 µg VSV-G using PEI. 48 hours post transfection, supernatants were pooled and concentrated using Amicon Ultra Centrifugal Filters. Concentrated lentiviral preps were stored at -80°C and titrated on A549 cells based on puromycin resistance. Briefly, 10-fold serial dilutions (from 10^−1^ to 10^−6^) were used to transduce 40,000 A549 cells in a 24 well plate format. 48 hours post transduction, cells were trypsinized and moved up to 6 well plates in the presence of 1.25 µg/mL puromycin. 9 days post transduction, cells were fixed, stained with Crystal Violet, and stained foci were counted to measure the number of cells surviving selection (i.e. those that were transduced with lentiCRISPRv2 harboring a puromycin resistance cassette). To perform the screen, 1.3 × 10^8^ A549 cells were transduced with lentiCRISPRv2-Brunello at an MOI of 0.3 in order to generate a population of single KO cells at high (>500X) coverage. Two days post transduction, cells were placed in selection with 1.25 µg/mL puromycin and passaged for 7 days, until there were no untransduced cells remaining. Thereafter, in triplicates with 8×10^6^ cells per flask, A549-Brunello cells were infected or not with HCoV-OC43 at an MOI of 0.1, and passaged for 7 days until >95% infection-induced cell death occurred. Cellular gDNA was isolated using Zymogen Quick-DNA Midiprep Plus Kit (Zymo Research), and sequencing libraries prepared via amplication of sgRNA loci utilizing F primers containing P5 and Read 1 Sequencing Primer and a R primer containing P7, a barcode, and the multiplexing Index Read Sequencing Primer, as described in Joung *et al*, 2017. Resulting libraries were gel purified, pooled, and sequenced on the Illumina HiSeq at Genewiz using 80 cycles of Read 1 (Forward) and 8 Cycles of Index 1 using standard Illumina sequencing primers.

### Pathway Analysis of screen hits

All 34 candidate genes were searched using the STRING database (https://string-db.org) for functional enrichment of protein-protein interactions using default settings, except the minimum required interaction score was changed from medium confidence (0.400) to high confidence (0.700). Subsequently, genes were annotated with UniProt keywords (https://uniprot.org)

### Validation of CRISPR hits

Individual sgRNAs targeting hits of interest were cloned into lentiCRISPRv2 via linearization with BsmBI followed by ligation of annealed oligos with compatible sticky ends using primers: VPS29 F: caccgGGACATCAAGTTATTCCATG: VPS29 R: aaacCATGGAATAACTTGATGTCCc; CCDC22 F: caccgCCGCAGGGTTGATCACACGC; CCDC22 R: aaacGCGTGTGATCAACCCTGCGGc; CCDC93 F: caccgTAGAATCCAAAGCTGATCCA; CCDC93 R: aaacTGGATCAGCTTTGGATTCTAc; COMMD3 F: caccgCTTGAAACATATCGACCCAG; COMMD3 R: aaacCTGGGTCGATATGTTTCAAGc. As a control, unmodified (“empty vector”) lentiCRISPRv2, which does not harbor an sgRNA cassette, was used. Lentiviral preparations were obtained by transfecting 1×10^6^ 293Ts with 1µg lentiCRISPRv2, 1µg NL-gagpol, and .2µg VSV-G using PEI. 2 days post transfection, supernatants were collected, filtered, and used to transduce 5×10^4^ A549-ACE2, HT1080-ACE2, 293T-ACE2, or NHBE cells. 2 days post transduction, cells were trypsinized, placed in selection with 1.25µg/mL puromycin, and passaged until there were no remaining viable untransduced cells. CRISPR KO was verified by Sanger Sequencing and Western Blot.

### Infectivity Assays

A total of 1×10^4^ cells per well were seeded on a 96-well plate in triplicate. The next day, cells were infected with each virus at an MOI of ∼0.3. For HCoV-OC43, HCoV-NL63, and HCoV-229E, infected plates were incubated at 34°C for 24 hours. For IAV, RSV, and adenovirus, infected plates were incubated at 37°C for 24 hours. For rVSV/SARS-CoV-2, infected plates were incubated at 37°C for 16 hours. Cells were then fixed in 4% PFA. For rVSV/SARS-CoV-2 and RSV, which encode eGFP and mKate2 reporter genes respectively, number of infected cells were measured directly by flow cytometry. Otherwise, cells were immunostained for viral antigens. Briefly, cells were blocked with 5.0% fetal bovine serum in PBS and permeabilized with 0.5% Saponin before a 30 minutes incubation with: HCoV-OC43: Anti-Coronavirus Group Antigen Antibody, nucleoprotein of OC-43 (1:1000, Sigma MAB9013); HCoV-NL63: Anti coronavirus NL63 (1:1000, Eurofins M.30.HCo.B2D4); HCoV-229: Anti coronavirus 229E (1:1000, Eurofins M.30.HCo.B1E7); IAV: Influenza A NP Antibody, FITC (1:50, Invitrogen MA1-7322); Adenovirus: Adenovirus Hexon Antibody, FITC (1:50, Invitrogen MA1-7329). For unconjugated primary antibodies (HCoV-OC43, HCoV-NL63, and HCoV-229E), a secondary antibody conjugate AF-488 Goat anti-Mouse IgG (H+L) (1:1000, Thermo) was used before infected cells were enumerated via flow cytometry.

### siRNA screening of VPS29 interactors

A list of well-known VPS29 interactors (Baños-Maetos, 2019 (Figure 1)) was selected and used to construct a targeted siRNA library constructed of a pool of four different gene specific siRNA sequences (ON TARGETplus SMARTpool siRNA, Dharmacon). siRNAs were reverse-transfected with 5×10^3^ HT1080-ACE2 using RNAiMAX (Thermo Scientific) according to the manufacturer’s protocol. Two-or three-days post transfection, cells were infected at an MOI of ∼0.3 and processed as above.

### Plasmid Construction

The lentiviral expression vector CSIN was derived from CSIB (Kane et al., 2013) by exchanging the Blasticidin resistance cassette with Neomycin. Briefly, primers Neo_CSIB_F: AAAAACACGATGATAATATGGCCACAACCAATTGAACAAGATGGATTGCACGCAGG TTCT and Neo_CSIB_R: AGCTTGATATCAAGCTTGCATGCCTGCAGGTCAGAAGAACTCGTCAAGAAGGCGAT AGAA were used to amplify the Neomycin resistance cassette and assemble into CSIB linearized with and BstXI and SbfI using NEBuilder HiFi DNA Assembly (NEB). The 2xFYVE-mSCAR endosome labeling construct was constructed by adding 2 FYVE domains to the N-terminus of mScarlett. FYVE domains were PCR amplified from the Hrs protein using primers FYVE_1_F: ACAGACTGAGTCGCCCGGGGGGGATCCGGCCGAGAGGGCCGCCACCGAGAGCGATGCCATGTTTGC, FYVE_1_R: GGCAGCAAACATGGCATCGCTCTCGGATCCTCCTCCTCCCTCCGCTTTCCTGTTCAGCTG, FYVE_2_F: CAGCTGAACAGGAAAGCGGAGGGAGGAGGAGGATCCGAGAGCGATGCCATGTTTG CTGCC, FYVE_2_R: TCACTGCCTCGCCCTTGCTCACCATGGATCCTCCTCCTCCCTCCGCTTTCCTGTTCAGCT. mScarlett was PCR amplified using primers mSCAR_F: AGCTGAACAGGAAAGCGGAGGGAGGAGGAGGATCCATGGTGAGCAAGGGCGAGGC AGTGA and mSCAR_R: GGGGGAGGGAGAGGGGCGGATCAGGCCAGAGAGGCCCTACTTGTACAGCTCGTCCA TGCC. The resulting fragments were assembled into CSIB linearized with SfiI using NEBuilder HiFi DNA Assembly (NEB).

To generate rVSV/SARS-CoV-2_NG-P_, The spike CDS was reverse transcribed and PCR amplified from rVSV/SARS-2/GFP_2E1_ using primers: P-NG-2E1-S-MluI_F: MLUAGAGATCGATCTGTTTCCTTGACACGCGTATGTTTGTGTTCCTGGTGCTGCTGCC A and P-NG-2E1-S-NotI_R: AACATGAAGAATCTGTTGTGCAGGGCGGCCGCCTTACAACAGGAGCCACAGGAA.

The resulting fragment was ligated into VSV NG-P (Jia et al., 2020) plasmid linearized with MluI and NotI using T4 ligase (NEB).

### Reconstitution experiments

A VPS29 coding sequence containing silent mutations in the sgRNA targeting sequence was purchased from IDT and cloned into CSIN using NEBuilder HiFi DNA Assembly (NEB). The VPS29_I91D_ and VPS29_L152E_ derivates were obtained via PCR mutagenesis using primers I91D F: GGTCACCAAGTAGATCCTTGGGGA, I91D R: TCCCCAAGGATCTACTTGGTGACC, L152E F: CCATCATTTGTGGAGATGGATATCCAGGC, L152E R: GCCTGGATATCCATCTCCACAAATGATGG. Resulting constructs, including an empty vector CSIN used as a control, were used to transduce single cell clones obtained from bulk EV or VPS29 KO HT1080-ACE2 via limiting dilution. Infectivity assays on the resulting cell lines were performed as above.

### HIV/Nanoluc CoV Pseudotype Assays

To generate HIV/Nanoluc CoV pseudotyped particles, 5×10^6^ 293T cells were plated in 10mL growth medium in a 10-cm dish. The next day, 7.5 µg pHIV-1_NL4-3_ ΔEnv-NanoLuc and 2.5 µg indicated CoV spike plasmid were transfected using PEI. Media was changed after 8 hours of incubation. After 48 hours post transfection, supernatant was harvested, passed through a 0.22-µm polyvinylidene fluoride syringe filter (Millipore; SLGVR33RS), aliquoted, and stored at - 80°C. To perform nanoluc assays with the resulting HIV/Nanoluc CoV pseudotyped particles, a total of 1×10^4^ HT1080-ACE2 WT or VPS29 KO cells per well were plated in triplicate in a 96-well plate. The next day, ∼1×10^3^ infectious units of HIV/Nanoluc CoV pseudotyped particles were added to cells and incubated at 37°C for 48 hours. Thereafter, cells were harvested for Nanoluc luciferase assays using the Nano-Glo® Luciferase Assay System (Promega, Cat# N1150).

### pHrodo Dextran Endocytosis assay

Cells were plated in a Nunc Lab-Tek II Chamber Slide (Thermo) at 5×10^3^ cells per well. The next day, cells were transduced with 2xFYVE-mSCAR to label endosomes. 48 hours post transduction, cells were treated with pHrodo Green Dextran 10,000 MW (Thermo, cat# P35368) at a concentration of 100 µg/mL for 60 minutes. Alternatively, unlabeled cells were treated with an equal ratio of pHrodo Red Dextran 10,000 MW (Thermo, cat# P10361) and AF-488 Dextran 10,000 MW (Thermo, cat# D22910). Thereafter, cells were washed 3X in PBS and placed in Live Cell Imaging Solution (Thermo Cat# A14291DJ). For 2x-FYVE labeled cells, images were acquired on a DeltaVision OMX SR imaging system using a 60X Widefield oil immersion objective (Olympus) with an exposure time of 50ms, 5.0% Transmission for the AF-488 channel, an exposure time of 50ms, 10% Transmission for the A568 channel, and an exposure time of 150ms, 10% Transmission for the DAPI channel. For co-Dextran-treated cells, images were acquired on a DeltaVision OMX SR imaging system using a 60X Widefield oil immersion objective (Olympus) with an exposure time of 25ms, 10.0% Transmission for the AF-488 channel, an exposure time of 50ms, 10% Transmission for the A568 channel, and an exposure time of 200ms, 10% Transmission for the DAPI channel.

### Microscopy of rVSV/SARS-CoV-2 infected cells

Cells were plated in a Nunc Lab-Tek II Chamber Slide (Thermo) at 5×10^3^ cells per well. The next day, cells were transduced with 2xFYVE-mSCAR to label endosomes. For rVSV/SARS-CoV-2_NG-P_ , 48 hours post transduction cells were treated with 5µM E64d (Sigma Aldrich E8640-250UG) for 30 minutes, followed by inoculation with rVSV/SARS-CoV-2_NG-P_ at an MOI of 2. 60 minutes post infection, cells were washed 3x with PBS and fixed in 4% PFA. Alternatively, unlabeled cells were treated with pHrodo Red Dextran and infected with rVSV/SARS-CoV-2_NG-P_ for 60 minutes. 60 minutes post infection, cells were washed 3X with PBS and imaged in Live Cell Imaging Solution. For cells with 2x-FYVE labeled endosomes, images were acquired on a DeltaVision OMX SR imaging system using a 60X Widefield oil immersion objective (Olympus) with an exposure time of 50ms, 10% Transmission for the AF-488 channel, an exposure time of 100ms, 10% Transmission for the A568 channel, and an exposure time of 150ms, 10% Transmission for the DAPI channel. For cells with Dextran Red labeled endosomes, images were acquired on a DeltaVision OMX SR imaging system using a 60X Widefield oil immersion objective (Olympus) with an exposure time of 50ms, 10% Transmission for the AF-488 channel, an exposure time of 50ms, 10% Transmission for the A568 channel, and an exposure time of 200ms, 10% Transmission for the DAPI channel.

### Influenza virus immunofluorescence

Cells were plated in a Nunc Lab-Tek II Chamber Slide (Thermo) at 5×10^3^ cells per well. The next day, cells were transduced with a construct expressing 2xFYVE-mSCAR to label endosomes. 48 hours post transduction, cells were infected with IAV at an MOI of ∼10. 60 minutes post infection, cells were washed with PBS, fixed in 4% PFA, permeabilized with 0.1% triton, blocked with FBS and stained for Influenza Virus Nucleoprotein (1:200, abcam cat# ab128193) and antibody conjugate AF-488 Goat anti-Mouse IgG (H+L) (Thermo, 1:1000). Images were acquired on a DeltaVision OMX SR imaging system using a 60X Widefield oil immersion objective (Olympus) with an exposure time of 50ms, 5.0% Transmission for the AF-488 channel, an exposure time of 100ms, 10% Transmission for the A568 channel, and an exposure time of 100ms, 10% Transmission for the DAPI channel.

### Quantification of fluorescence microscopy

For each cell, Regions Of Interest (ROIs) corresponding to labeled endosomes were defined using the freehand selection tool in Fiji. Quantification of mean fluorescence intensity inside each ROI was determined using the Measure command. For punctae quantification, the number of punctae inside each ROI was counted and summed to give the total number of punctae inside ROIs for each cell. Additionally, the total number of punctae outside of ROIs in each cell was measured. The reported % of virus in endosomes corresponds to:

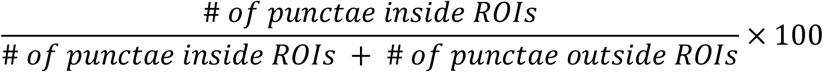

### Cathepsin L activity assay

Cells were plated 2×10^4^ cells per well in a Nunc Lab-Tek II Chamber Slide (Thermo). The next day, intracellular cathepsin L activity was detected using the Magic Red Cathepsin L Assay Kit (Biorad cat# ICT941). Briefly, cells were incubated in 1X Magic Red and Hoechst 33342 Stain for 30 minutes, then washed 3X with PBS before being placed in Live Cell Imaging Solution (Thermo Cat# A14291DJ). Images were acquired on a DeltaVision OMX SR imaging system using a 60X Widefield oil immersion objective (Olympus) using an exposure time of 50ms, 10% Transmission for the A568 nm channel and an exposure time of 100ms, 10% Transmission for the DAPI channel.

### Cathepsin L localization staining

The coding sequence of CTSL was tagged with a 3’V5 and cloned into CSIN using NEBuilder HiFi DNA Assembly (NEB) with primers CTSL_3’_V5_F: ACAGACTGAGTCGCCCGGGGGGGATCCGGCCGAGAGGGCCGCCACCATGAATCCTACACTCATCCTTGC and CTSL_3’_V5_R: GGGGGAGGGAGAGGGGCGGATCAGGCCAGAGAGGCCTCACGTAGAATCGAGACCGAGGAGAGGGTTAGGGATAGGCTTACCCACAGTGGGGTAGCTGGCT. Cells stably expressing this 3’V5-tagged CTSL were plated in a Nunc Lab-Tek II Chamber Slide (Thermo) at 5×10^3^ cells per well. The next day, cells were transduced with a construct expressing 2xFYVE-mSCAR to label endosomes. 48 hours post transduction, cells were fixed in 4% PFA, permeabilized with 0.1% triton, blocked with FBS and stained for V5 (invitrogen cat# 46-0705, 1:1000) and antibody conjugate AF-488 Goat anti-Mouse IgG (H+L) (Thermo, 1:1000). Images were acquired on a DeltaVision OMX SR imaging system using a 60X Widefield oil immersion objective (Olympus) with an exposure time of 50ms, 5.0% Transmission for the AF-488 channel, an exposure time of 100ms, 10% Transmission for the A568 channel, and an exposure time of 100ms, 10% Transmission for the DAPI channel.

## QUANTIFICATION AND STATISTICAL ANALYSIS

Raw FASTQ files were aligned to the Brunello library and scored using the MAGeCK statistical package. All flow cytometry data were analyzed using FlowJo software, version 10.6.1. All graphs were generated using GraphPad Prism, version 8. Error bars correspond to the standard deviation. All images were generated by maximum intensity projection using Fiji (https://fiji.sc/).

## SUPPLEMENTAL INFORMATION

**Supplemental Figure S1:**
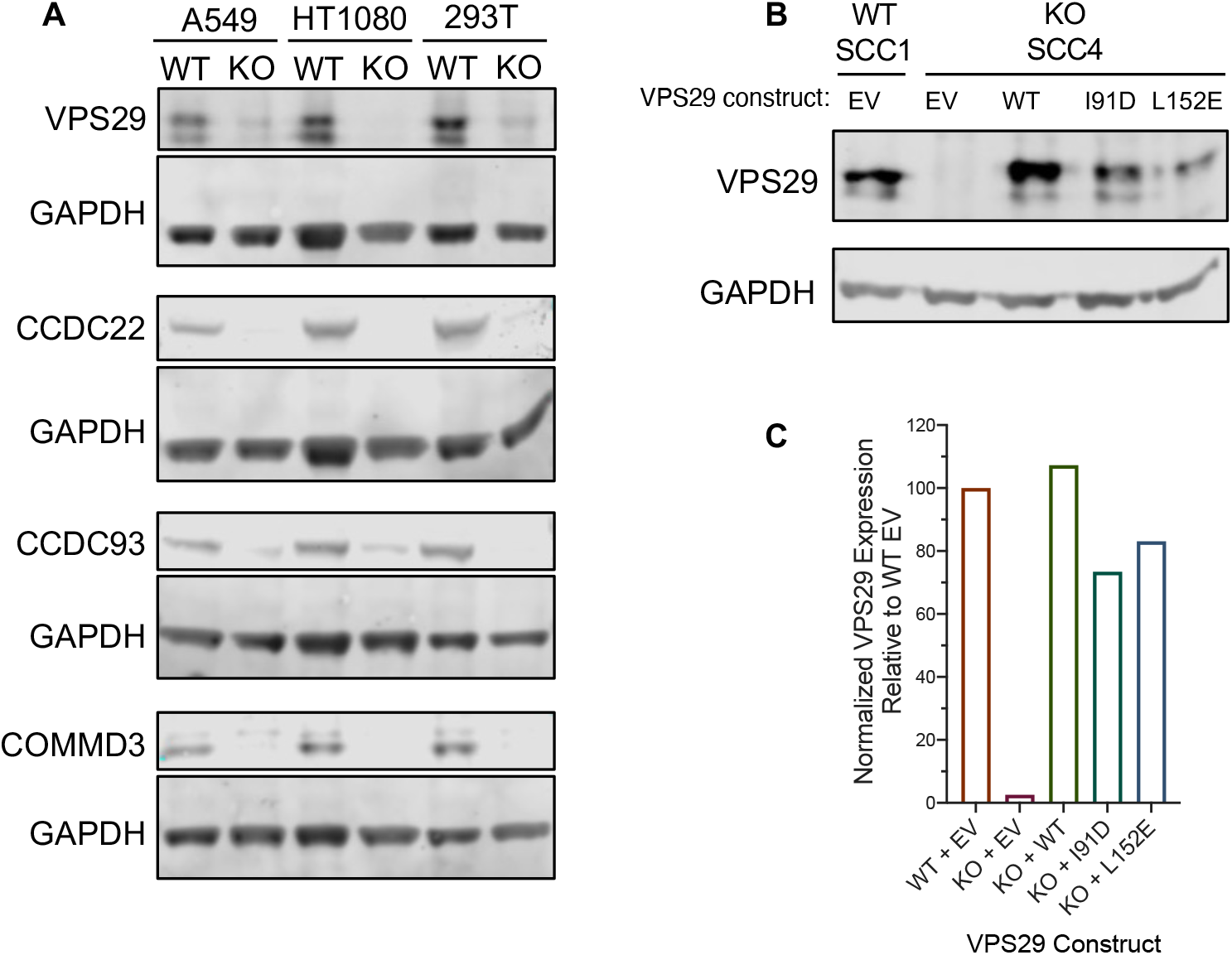
Validation of CRISPR KO and VPS29 Reconstitution. (A): Western Blot confirmation of VPS29, CCDC22, CCDC93, and COMMD3 KO. (B): Western Blot confirmation of VPS29 SSC KO and reconstitution. Antibodies: VPS29 (ab236796), CCDC22 (protein tech 16636-1-AP), CCDC93 (protein tech 20861-1-AP), COMMD3 (protein tech 26240-1-AP). (C): Quantification of the level of VPS29 expression normalized to GAPDH.

**Supplemental Figure S2:**
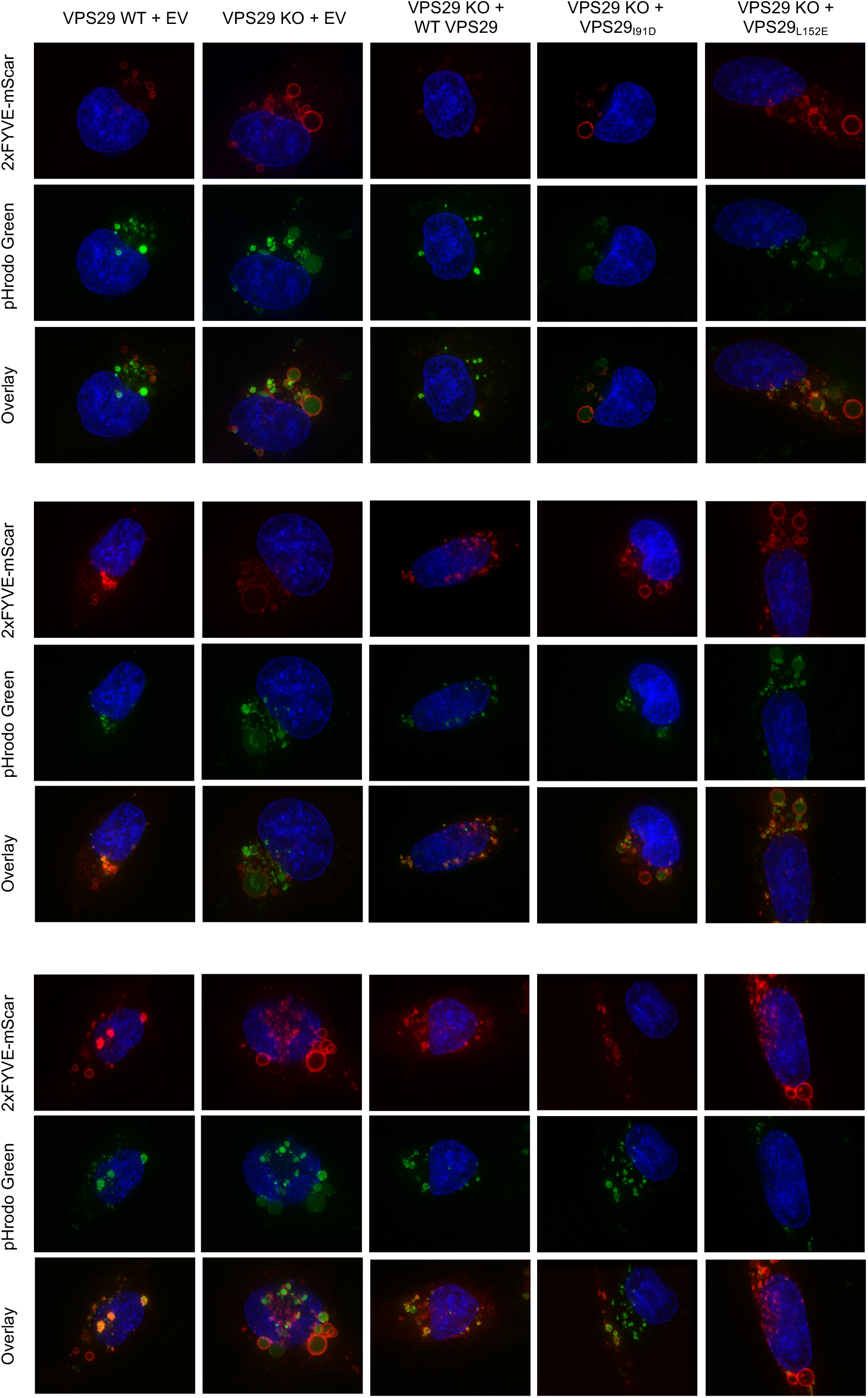
Additional representative images demonstrating VPS29 KO results in enlarged, deacidified PI(3)P-rich vesicles. Additional representative images from Figure 4. HT1080 cells transduced with 2xFYVE-mSCAR after incubation with pHrodo Green Dextran for 60 minutes

**Supplemental Figure S3:**
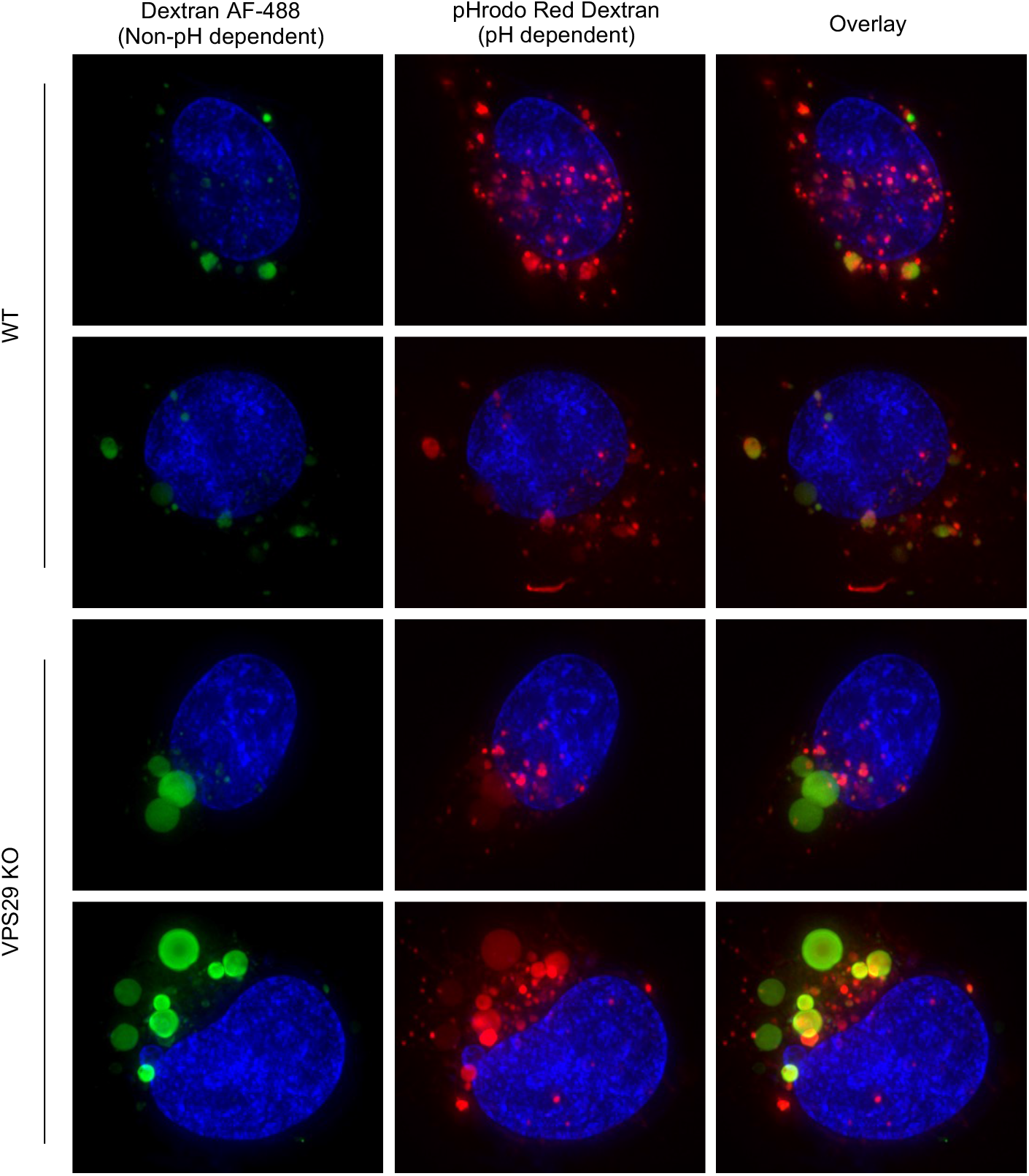
Additional representative images showing the enlarged, deacidified vesicles in VPS29-KO cells are not impaired for cargo loading. Additional representative images from Figure 5. WT and VPS29 KO HT1080 cells incubated for 60 minutes with an equal molar ratio of pHrodo Dextran Red 10,000 MW and Dextran AF-488 10,000 MW.

**Supplemental Figure S4:**
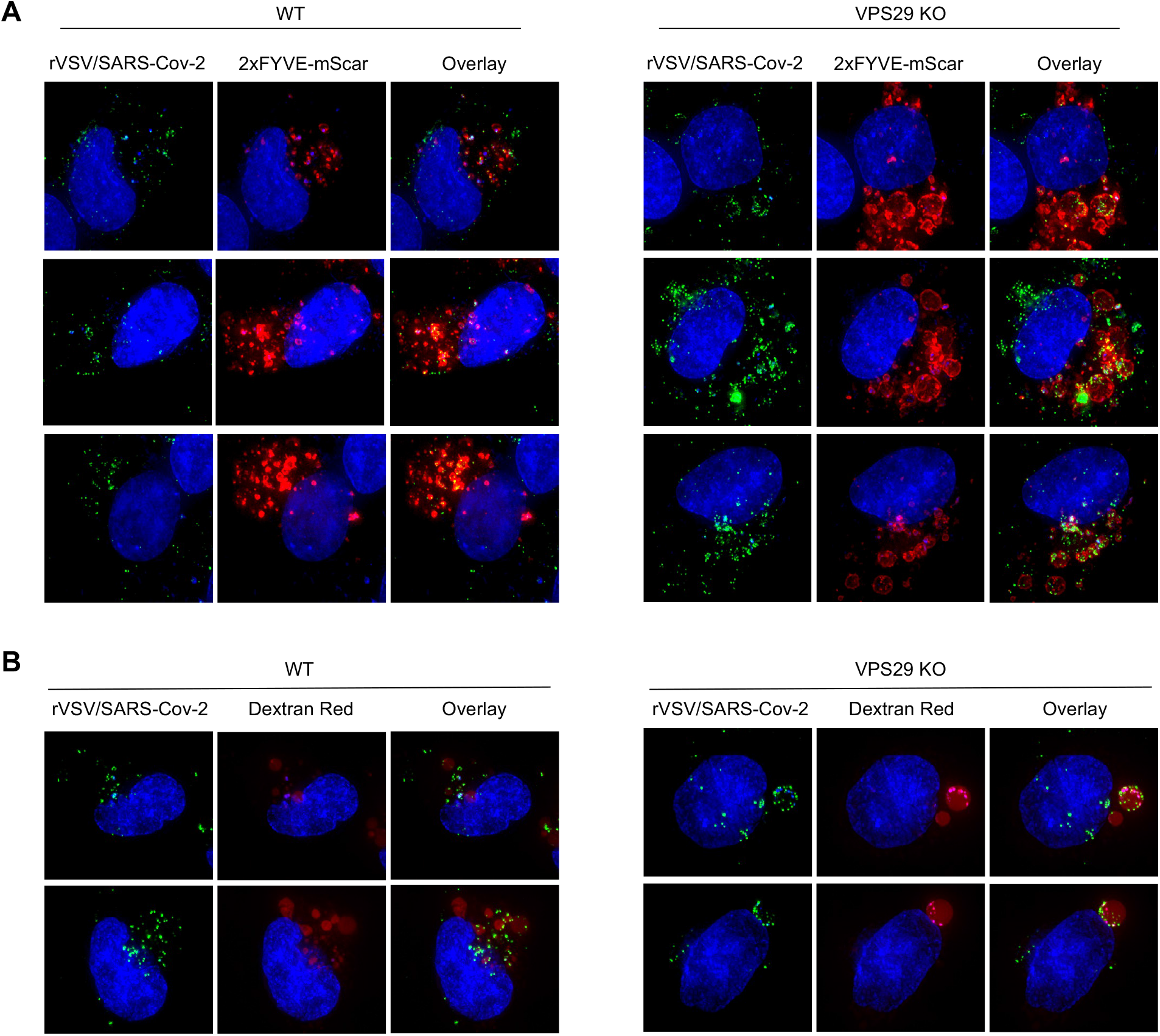
Additional representative images demonstrating VPS29 KO results in rVSV/SARS-CoV-2 remaining trapped in vesicles. (A): Additional representative images from Figure 6A. rVSV/SARS-CoV-2_NG-P_ infection in WT and VPS29 KO HT1080 cells. Cells were infected with rVSV/SARS-CoV-2_NG-P_ for 60 minutes. (B): Additional representative images from Figure 6B. WT and VPS29 KO HT1080 cells incubated for 60 minutes with Dextran Red 10,000 MW and rVSV/SARS-CoV-2_NG-P._

**Supplemental Figure S5:**
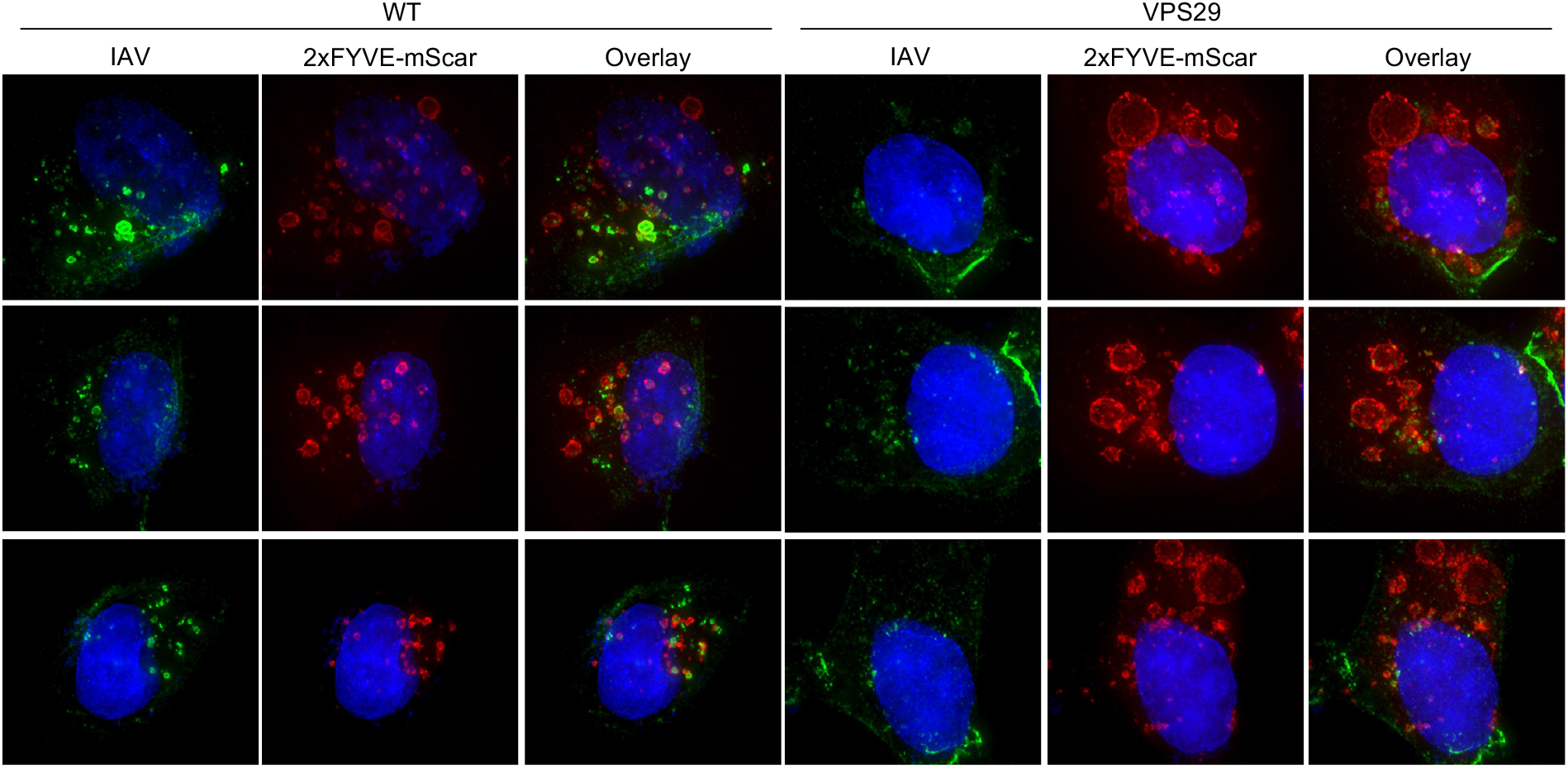
Additional representative images demonstrating that IAV is more associated with PI(3)P-rich endosomal membranes in WT HT1080 cells than in VPS29 KO HT1080 cells. Additional representative images from Figure 6C: IAV infection in WT and VPS29 KO HT1080 cells labeled with 2xFYVE-mSCAR. Cells were infected with IAV for 60 minutes then fixed and stained for IAV NP.

**Supplemental Figure S6:**
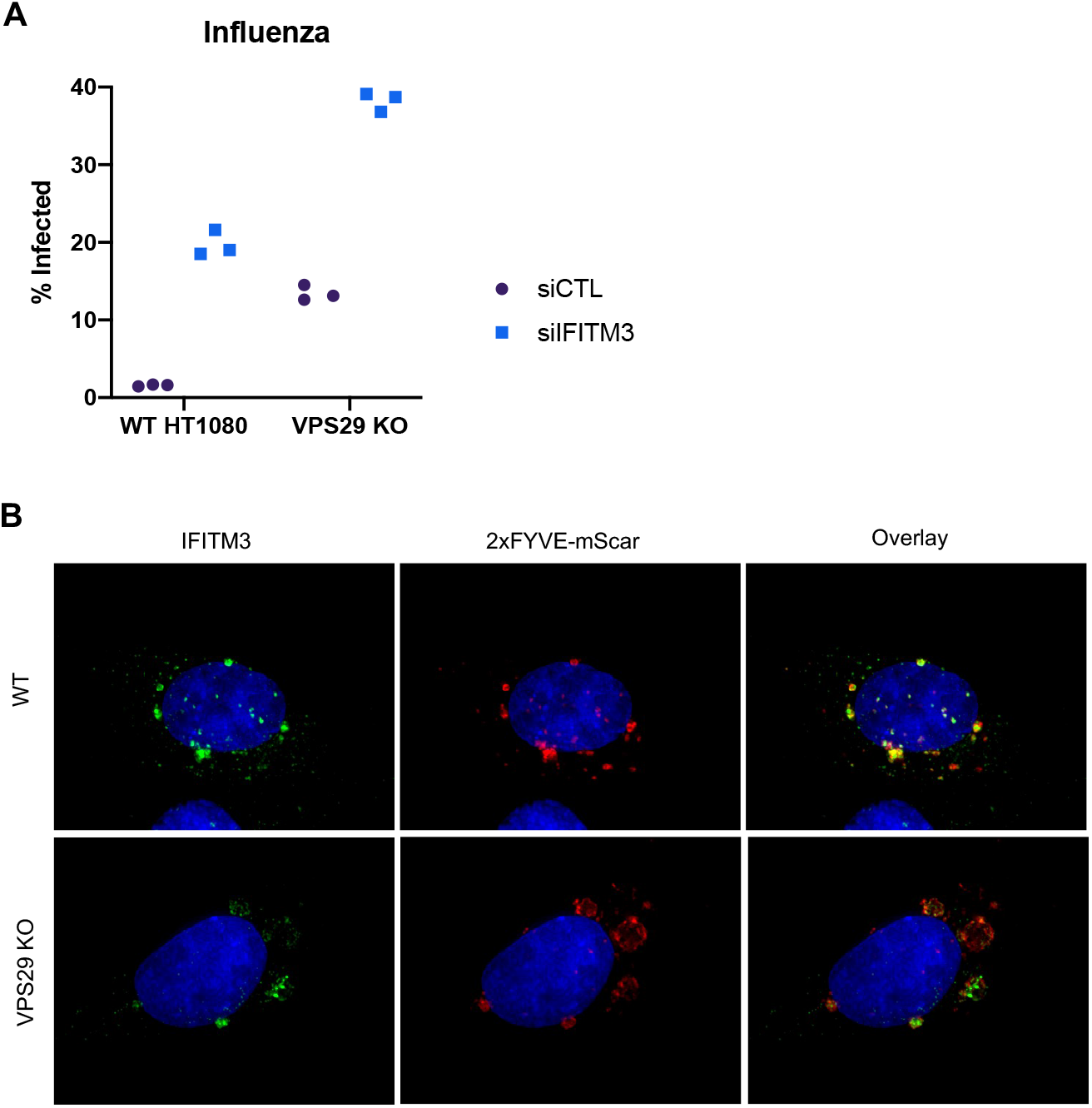
The enhancement of Influenza infection by VPS29 KO is not mediated by loss of IFITM3 activity. (A): WT and VPS29 KO HT1080 cells were transfected with a pool of four control siRNAs (siCTL) or a pool of four siRNAs targeting IFITM3 (siIFITM3). Three days post transfection, cells were infected with IAV. At 24 hours post infection, cells were stained, and the percent infected cells was determined by flow cytometry. (B): Representative images of WT and VPS29 KO HT1080 cells stably expressing V5-tagged IFITM3 and labeled with 2xFYVE-mSCAR.

**Supplemental Figure S7:**
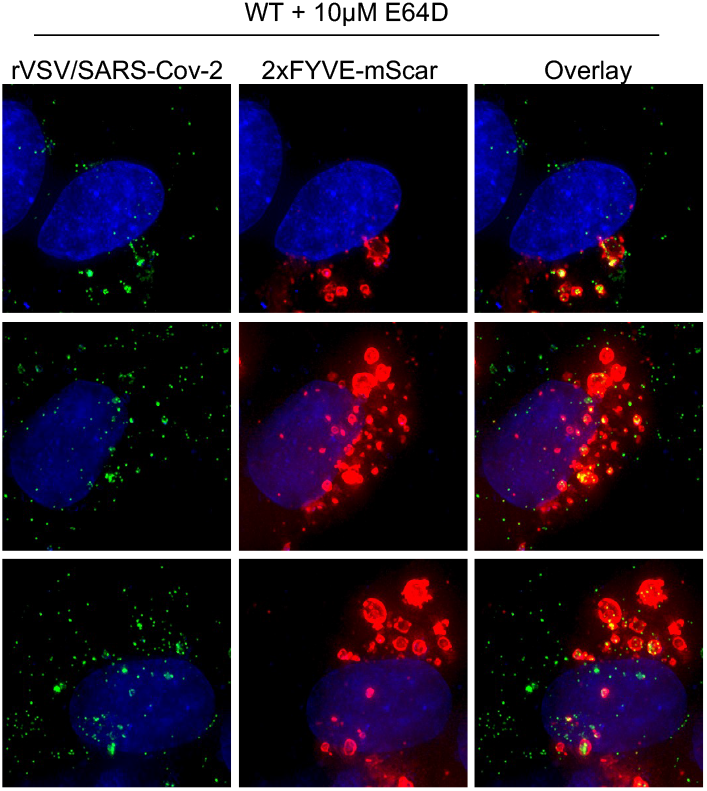
Additional representative images demonstrating cathepsin inhibition with E64d phenocopies VPS29 KO. Additional representative images from Figure 7I: 2xFYVE-mSCAR labeled cells were treated with E64d for 30 minutes, then infected with rVSV/SARS-CoV-2_NG-P_ for 60 minutes.

## Bibliography

Baños-Mateos, S., Rojas, A.L., and Hierro, A. (2019). VPS29, a tweak tool of endosomal recycling. Curr. Opin. Cell Biol. 59, 81–87.

Böttcher-Friebertshäuser, E., Garten, W., Matrosovich, M., and Klenk, H.D. (2014). The hemagglutinin: a determinant of pathogenicity. Curr. Top. Microbiol. Immunol. 385, 3–34.

Burda, P., Padilla, S.M., Sarkar, S., and Emr, S.D. (2002). Retromer function in endosome-to-Golgi retrograde transport is regulated by the yeast Vps34 PtdIns 3-kinase. J. Cell Sci. 115, 3889–3900.

Collins, B.M., Skinner, C.F., Watson, P.J., Seaman, M.N.J., and Owen, D.J. (2005). Vps29 has a phosphoesterase fold that acts as a protein interaction scaffold for retromer assembly. Nat. Struct. Mol. Biol. 12, 594–602.

Coronaviridae Study Group of the International Committee on Taxonomy of Viruses (2020). The species Severe acute respiratory syndrome-related coronavirus: classifying 2019-nCoV and naming it SARS-CoV-2. Nat. Microbiol. 5, 536–544.

Coutard, B., Valle, C., de Lamballerie, X., Canard, B., Seidah, N.G., and Decroly, E. (2020). The spike glycoprotein of the new coronavirus 2019-nCoV contains a furin-like cleavage site absent in CoV of the same clade. Antiviral Res. 176, 104742.

Cui, Y., Carosi, J.M., Yang, Z., Ariotti, N., Kerr, M.C., Parton, R.G., Sargeant, T.J., and Teasdale, R.D. (2019). Retromer has a selective function in cargo sorting via endosome transport carriers. J. Cell Biol. 218, 615–631.

Daniloski, Z., Jordan, T.X., Wessels, H.H., Hoagland, D.A., Kasela, S., Legut, M., Maniatis, S., Mimitou, E.P., Lu, L., Geller, E., et al. (2020). Identification of Required Host Factors for SARS-CoV-2 Infection in Human Cells. Cell.

Del Olmo, T., Lauzier, A., Normandin, C., Larcher, R., Lecours, M., Jean, D., Lessard, L., Steinberg, F., Boisvert, F.-M., and Jean, S. (2019). APEX2-mediated RAB proximity labeling identifies a role for RAB21 in clathrin-independent cargo sorting. EMBO Rep. 20.

Doench, J.G., Fusi, N., Sullender, M., Hegde, M., Vaimberg, E.W., Donovan, K.F., Smith, I., Tothova, Z., Wilen, C., Orchard, R., et al. (2016). Optimized sgRNA design to maximize activity and minimize off-target effects of CRISPR-Cas9. Nat. Biotechnol. 34, 184–191.

Feeley, E.M., Sims, J.S., John, S.P., Chin, C.R., Pertel, T., Chen, L.-M., Gaiha, G.D., Ryan, B.J., Donis, R.O., Elledge, S.J., et al. (2011). IFITM3 inhibits influenza A virus infection by preventing cytosolic entry. PLoS Pathog. 7, e1002337.

Gao, H., Lin, Y., He, J., Zhou, S., Liang, M., Huang, C., Li, X., Liu, C., and Zhang, P. (2019). Role of heparan sulfate in the Zika virus entry, replication, and cell death. Virology 529, 91–100.

Gillooly, D.J., Morrow, I.C., Lindsay, M., Gould, R., Bryant, N.J., Gaullier, J.M., Parton, R.G., and Stenmark, H. (2000). Localization of phosphatidylinositol 3-phosphate in yeast and mammalian cells. EMBO J. 19, 4577–4588.

Grove, J., and Marsh, M. (2011). The cell biology of receptor-mediated virus entry. J. Cell Biol. 195, 1071–1082.

Harbour, M.E., Breusegem, S.Y.A., Antrobus, R., Freeman, C., Reid, E., and Seaman, M.N.J. (2010). The cargo-selective retromer complex is a recruiting hub for protein complexes that regulate endosomal tubule dynamics. J. Cell Sci. 123, 3703–3717.

Hoffmann, H.-H., Sánchez-Rivera, F.J., Schneider, W.M., Luna, J.M., Soto-Feliciano, Y.M., Ashbrook, A.W., Le Pen, J., Leal, A.A., Ricardo-Lax, I., Michailidis, E., et al. (2021). Functional interrogation of a SARS-CoV-2 host protein interactome identifies unique and shared coronavirus host factors. Cell Host Microbe 29, 267–280.e5.

Hotard, A.L., Shaikh, F.Y., Lee, S., Yan, D., Teng, M.N., Plemper, R.K., Crowe, J.E., and Moore, M.L. (2012). A stabilized respiratory syncytial virus reverse genetics system amenable to recombination-mediated mutagenesis. Virology 434, 129–136.

Jerala, R., Zerovnik, E., Kidric, J., and Turk, V. (1998). pH-induced conformational transitions of the propeptide of human cathepsin L. A role for a molten globule state in zymogen activation. J. Biol. Chem. 273, 11498–11504.

Jia, D., Zhang, J.-S., Li, F., Wang, J., Deng, Z., White, M.A., Osborne, D.G., Phillips-Krawczak, C., Gomez, T.S., Li, H., et al. (2016). Structural and mechanistic insights into regulation of the retromer coat by TBC1d5. Nat. Commun. 7, 13305.

Jia, M., Liberatore, R.A., Guo, Y., Chan, K.-W., Pan, R., Lu, H., Waltari, E., Mittler, E., Chandran, K., Finzi, A., et al. (2020). VSV-Displayed HIV-1 Envelope Identifies Broadly Neutralizing Antibodies Class-Switched to IgG and IgA. Cell Host Microbe 27, 963–975.e5.

Joung, J., Konermann, S., Gootenberg, J.S., Abudayyeh, O.O., Platt, R.J., Brigham, M.D., Sanjana, N.E., and Zhang, F. (2017). Genome-scale CRISPR-Cas9 knockout and transcriptional activation screening. Nat. Protoc. 12, 828–863.

Kane, M., Yadav, S.S., Bitzegeio, J., Kutluay, S.B., Zang, T., Wilson, S.J., Schoggins, J.W., Rice, C.M., Yamashita, M., Hatziioannou, T., et al. (2013). MX2 is an interferon-induced inhibitor of HIV-1 infection. Nature 502, 563–566.

Killerby, M.E., Biggs, H.M., Haynes, A., Dahl, R.M., Mustaquim, D., Gerber, S.I., and Watson, J.T. (2018). Human coronavirus circulation in the United States 2014-2017. J. Clin. Virol. 101, 52–56.

Krzyzaniak, M.A., Zumstein, M.T., Gerez, J.A., Picotti, P., and Helenius, A. (2013). Host cell entry of respiratory syncytial virus involves macropinocytosis followed by proteolytic activation of the F protein. PLoS Pathog. 9, e1003309.

Lakadamyali, M., Rust, M.J., and Zhuang, X. (2004). Endocytosis of influenza viruses. Microbes Infect. 6, 929–936.

Laporte, M., and Naesens, L. (2017). Airway proteases: an emerging drug target for influenza and other respiratory virus infections. Curr Opin Virol 24, 16–24.

Li, W., Xu, H., Xiao, T., Cong, L., Love, M.I., Zhang, F., Irizarry, R.A., Liu, J.S., Brown, M., and Liu, X.S. (2014). MAGeCK enables robust identification of essential genes from genome-scale CRISPR/Cas9 knockout screens. Genome Biol. 15, 554.

Luteijn, R.D., van Diemen, F., Blomen, V.A., Boer, I.G.J., Manikam Sadasivam, S., van Kuppevelt, T.H., Drexler, I., Brummelkamp, T.R., Lebbink, R.J., and Wiertz, E.J. (2019). A Genome-Wide Haploid Genetic Screen Identifies Heparan Sulfate-Associated Genes and the Macropinocytosis Modulator TMED10 as Factors Supporting Vaccinia Virus Infection. J. Virol. 93.

Marsh, M., and Helenius, A. (2006). Virus entry: open sesame. Cell 124, 729–740.

McNally, K.E., Faulkner, R., Steinberg, F., Gallon, M., Ghai, R., Pim, D., Langton, P., Pearson, N., Danson, C.M., Nägele, H., et al. (2017). Retriever is a multiprotein complex for retromer-independent endosomal cargo recycling. Nat. Cell Biol. 19, 1214–1225.

Meier, O., and Greber, U.F. (2004). Adenovirus endocytosis. J Gene Med 6 Suppl 1, S152–63.

Milewska, A., Zarebski, M., Nowak, P., Stozek, K., Potempa, J., and Pyrc, K. (2014). Human coronavirus NL63 utilizes heparan sulfate proteoglycans for attachment to target cells. J. Virol. 88, 13221–13230.

Millet, J.K., and Whittaker, G.R. (2015). Host cell proteases: Critical determinants of coronavirus tropism and pathogenesis. Virus Res. 202, 120–134.

Mulherkar, N., Raaben, M., de la Torre, J.C., Whelan, S.P., and Chandran, K. (2011). The Ebola virus glycoprotein mediates entry via a non-classical dynamin-dependent macropinocytic pathway. Virology 419, 72–83.

Park, R.J., Wang, T., Koundakjian, D., Hultquist, J.F., Lamothe-Molina, P., Monel, B., Schumann, K., Yu, H., Krupzcak, K.M., Garcia-Beltran, W., et al. (2017). A genome-wide CRISPR screen identifies a restricted set of HIV host dependency factors. Nat. Genet. 49, 193– 203.

Phillips-Krawczak, C.A., Singla, A., Starokadomskyy, P., Deng, Z., Osborne, D.G., Li, H., Dick, C.J., Gomez, T.S., Koenecke, M., Zhang, J.-S., et al. (2015). COMMD1 is linked to the WASH complex and regulates endosomal trafficking of the copper transporter ATP7A. Mol. Biol. Cell 26, 91–103.

Popa, A., Zhang, W., Harrison, M.S., Goodner, K., Kazakov, T., Goodwin, E.C., Lipovsky, A., Burd, C.G., and DiMaio, D. (2015). Direct binding of retromer to human papillomavirus type 16 minor capsid protein L2 mediates endosome exit during viral infection. PLoS Pathog. 11, e1004699.

Rojas, R., van Vlijmen, T., Mardones, G.A., Prabhu, Y., Rojas, A.L., Mohammed, S., Heck, A.J.R., Raposo, G., van der Sluijs, P., and Bonifacino, J.S. (2008). Regulation of retromer recruitment to endosomes by sequential action of Rab5 and Rab7. J. Cell Biol. 183, 513–526.

Sanson, K.R., Hanna, R.E., Hegde, M., Donovan, K.F., Strand, C., Sullender, M.E., Vaimberg, E.W., Goodale, A., Root, D.E., Piccioni, F., et al. (2018). Optimized libraries for CRISPR-Cas9 genetic screens with multiple modalities. Nat. Commun. 9, 5416.

Schmidt, F., Weisblum, Y., Muecksch, F., Hoffmann, H.-H., Michailidis, E., Lorenzi, J.C.C., Mendoza, P., Rutkowska, M., Bednarski, E., Gaebler, C., et al. (2020). Measuring SARS-CoV-2 neutralizing antibody activity using pseudotyped and chimeric viruses. J. Exp. Med. 217.

Schneider, W.M., Luna, J.M., Hoffmann, H.-H., Sánchez-Rivera, F.J., Leal, A.A., Ashbrook, A.W., Le Pen, J., Ricardo-Lax, I., Michailidis, E., Peace, A., et al. (2020). Genome-scale identification of SARS-CoV-2 and pan-coronavirus host factor networks. Cell.

Schornberg, K., Matsuyama, S., Kabsch, K., Delos, S., Bouton, A., and White, J. (2006). Role of endosomal cathepsins in entry mediated by the Ebola virus glycoprotein. J. Virol. 80, 4174– 4178.

Schott, D.H., Cureton, D.K., Whelan, S.P., and Hunter, C.P. (2005). An antiviral role for the RNA interference machinery in Caenorhabditis elegans. Proc. Natl. Acad. Sci. USA 102, 18420– 18424.

Schwegmann-Wessels, C., and Herrler, G. (2006). Sialic acids as receptor determinants for coronaviruses. Glycoconj. J. 23, 51–58.

Seaman, M.N.J., Gautreau, A., and Billadeau, D.D. (2013). Retromer-mediated endosomal protein sorting: all WASHed up! Trends Cell Biol. 23, 522–528.

Singla, A., Fedoseienko, A., Giridharan, S.S.P., Overlee, B.L., Lopez, A., Jia, D., Song, J., Huff-Hardy, K., Weisman, L., Burstein, E., et al. (2019). Endosomal PI(3)P regulation by the COMMD/CCDC22/CCDC93 (CCC) complex controls membrane protein recycling. Nat. Commun. 10, 4271.

Szklarczyk, D., Gable, A.L., Lyon, D., Junge, A., Wyder, S., Huerta-Cepas, J., Simonovic, M., Doncheva, N.T., Morris, J.H., Bork, P., et al. (2019). STRING v11: protein-protein association networks with increased coverage, supporting functional discovery in genome-wide experimental datasets. Nucleic Acids Res. 47, D607–D613.

UniProt Consortium (2019). UniProt: a worldwide hub of protein knowledge. Nucleic Acids Res. 47, D506–D515.

Volchkov, V., and Klenk, H.D. (2018). Proteolytic processing of filovirus glycoproteins. In Activation of Viruses by Host Proteases,E. Böttcher-Friebertshäuser, W. Garten, and H.D. Klenk, eds. (Cham: Springer International Publishing), pp. 99–108.

Wang, R., Simoneau, C.R., Kulsuptrakul, J., Bouhaddou, M., Travisano, K.A., Hayashi, J.M., Carlson-Stevermer, J., Zengel, J.R., Richards, C.M., Fozouni, P., et al. (2021). Genetic Screens Identify Host Factors for SARS-CoV-2 and Common Cold Coronaviruses. Cell 184, 106–119.e14.

Whelan, S.P., Barr, J.N., and Wertz, G.W. (2000). Identification of a minimal size requirement for termination of vesicular stomatitis virus mRNA: implications for the mechanism of transcription. J. Virol. 74, 8268–8276.

White, J.M., and Whittaker, G.R. (2016). Fusion of enveloped viruses in endosomes. Traffic 17, 593–614.

Winstone, H., Lista, M.J., Reid, A.C., Bouton, C., Pickering, S., Galao, R.P., Kerridge, C., Doores, K.J., Swanson, C., and Neil, S. (2021). The polybasic cleavage site in the SARS-CoV-2 spike modulates viral sensitivity to Type I interferon and IFITM2. J. Virol.

Zhu, Y., Feng, F., Hu, G., Wang, Y., Yu, Y., Zhu, Y., Xu, W., Cai, X., Sun, Z., Han, W., et al. (2021). A genome-wide CRISPR screen identifies host factors that regulate SARS-CoV-2 entry. Nat. Commun. 12, 961.

